# Transplantation-based screen uncovers inducers of muscle progenitor cell engraftment across vertebrate species

**DOI:** 10.1101/2023.01.13.523942

**Authors:** Sahar Tavakoli, Isaac Adatto, Sara Ashrafi Kakhki, Victoria S Chan, Haleh Fotowat, Eric Gähwiler, Margot E Manning, Kathleen A Messemer, Apoorva Rangan, Song Yang, Amy J Wagers, Leonard I Zon

## Abstract

Stem cell transplantation presents a potentially curative strategy for genetic disorders of skeletal muscle, but this approach is limited due to the deleterious effects of cell expansion *in vitro* and consequent poor engraftment efficiency. In an effort to overcome this limitation, we sought to identify molecular signals that enhance the myogenic activity of cultured muscle progenitors. Here, we report the development and application of a cross-species small molecule screening platform employing zebrafish and mouse, which enables rapid, direct evaluation of the effects of chemical compounds on the engraftment of transplanted muscle precursor cells. Using this system, we screened a library of bioactive lipids to identify those that could increase myogenic engraftment *in vivo* in zebrafish and mice. Two lipids, lysophosphatidic acid (LPA) and niflumic acid (NFA), are linked to activation of intracellular calcium ion flux, which showed conserved, dose-dependent and synergistic effects in promoting muscle engraftment across these vertebrate species.

## Introduction

Skeletal muscle is subject to mechanical and physiological stresses throughout life and requires maintenance and repair, including replacement of myonuclei, to sustain its function. The basic structural unit of skeletal muscle is the myofiber, a multinucleated and post-mitotic cell (Huxley, 1963). Because myofibers are multinucleated and structurally complex, they cannot replace themselves during tissue repair through traditional cell division (Carlson, 1973; Okazaki and Holtzer, 1966). Instead, myofiber repair is contingent upon the successful fusion of mononucleated muscle progenitors, called myoblasts, into de novo or residual injured or atrophied fibers (Partridge et al., 1978).

Myoblasts are primarily generated from resident muscle stem cells, called “satellite cells,” which are Pax7^+^/MyoD^-^/Myf5^-^ mononuclear, unipotent stem cells (Bischoff, 1975; Conboy et al., 2003; Konigsberg et al., 1975; Sacco et al., 2008; Sherwood et al., 2004; Yin et al., 2013). Satellite cells localize beneath the basal lamina of muscle fibers (Mauro, 1961) and generally remain mitotically quiescent in intact (uninjured) adult muscle. Satellite cells undergo activation upon muscle damage, initiating both self-renewing and differentiating divisions and producing daughter myoblasts that fuse together to build, and re-build, new muscle (Collins et al., 2005). Chronic muscle damage, caused by repeated injury or genetic disorders like muscular dystrophies, may deplete satellite cells and impair their function, leading to a diminished regenerative response, progressive loss of muscle mass, and reduction of strength and mobility (Carlson and Faulkner, 1996; Conboy et al., 2003; Cox et al., 1993; Ervasti, 2007). Transplantation-based studies in animal models have demonstrated the utility of engrafted satellite cells for regenerating diseased muscle (Cerletti et al., 2008; Fukada et al., 2004; Kuang et al., 2007; Lean et al., 2019; Montarras et al., 2005; Sacco et al., 2008; Tanaka et al., 2009). However, challenges in obtaining adequate numbers of satellite cells from adult skeletal muscle and in achieving sufficient efficiencies for engraftment of culture-expanded cells, have presented significant barriers to the clinical application of such transplantation approaches (Tabebordbar et al., 2013).

To address these challenges, we devised a novel screening approach that leverages the ability to rapidly and cost-effectively assess the impact of defined chemical compounds on the engraftment efficiency of muscle progenitors *in vivo* in zebrafish. Potential pro-myogenic effects of compounds identified through this screen were subsequently validated in assays using highly purified mouse muscle satellite cells (Cerletti et al., 2008; Sherwood et al., 2004). The throughput and accessibility of this model allowed us to evaluate 187 compounds sourced from the ICCB Known Bioactives Library, focusing primarily on lipid mediators, a relatively understudied class of biomolecules that can play key roles in enhancing cell migration and promoting regenerative function (Li et al., 2015; Oh et al., 2016). Candidate compounds were added to the cells *in vitro*, and 4 hours later cells were washed and transplanted. This approach sought compounds that had specific effects on muscle engraftment, limiting the effect of the compounds to the donor cells with little if any exposure of the recipient muscle tissue. These efforts identified two small molecules, lysophosphatidic acid (LPA) and niflumic acid (NFA), both of which increased the engraftment efficiency and muscle-forming activity of transplanted myogenic progenitors in zebrafish and in mice. RNA sequencing and calcium ion imaging studies further revealed that both compounds alter the expression patterns of ion transport genes and increase intracellular calcium concentrations, suggesting that the effects of these lipids are mediated via regulation of calcium-dependent second messenger systems (Berridge et al., 2000). Finally, supporting the potential translational relevance of these compounds, we documented improved physiologic function in swimming performance tests of dystrophic zebrafish (Guyon et al., 2009) transplanted with NFA-treated or LPA-treated muscle precursors. In summary, the novel cross-species approach developed in this study has uncovered previously unknown inducers of skeletal muscle cell engraftment and suggests new potential opportunities for muscle cell therapy as a treatment option for degenerative muscle disorders.

## Results

### Establishing transplantation parameters for zebrafish muscle cells using a limiting dilution assay

Zebrafish myogenic progenitor cells (ZeMPCs) were generated by *in vitro* culture of blastomeres from *mylz2-*GFP, *mylz2-*mCherry or *myf5-*GFP*;mylz2-*mCherry double transgenic zebrafish embryos by adaptation of published protocols (Xu et al., 2013), with *myf5* serving as a marker of myogenic progenitors and *mylz2* serving as a marker of terminally differentiated muscle cells (Ju et al., 2003). ZeMPCs were transplanted into the flank muscles on both sides of transparent *casper* zebrafish recipients (**Figure 1A**). At 7 days post-transplantation (dpt), engraftment success was assessed by *in vivo* fluorescent stereo microscopy to detect GFP or mCherry-tagged donor-engrafted myofibers in the recipient fish. Notably, we observed persistence of donor-engrafted muscle cells in recipient fish for up to one-year post-transplantation (**Figure 1B and 1C**). The optimal number of transplanted cells for screening was determined by performing an extreme limiting dilution analysis (ELDA) (Hu and Smyth, 2009) for engraftment. These initial studies yielded a 100% successful engraftment rate in fish receiving 700 ZeMPCs, and a 30% successful engraftment rate in fish receiving 100-cell transplants **(Table S1)**. Using ELDA software (Hu and Smyth, 2009), we calculated the cell potency, which implies 1 out of 272 ZeMPCs produced under these conditions successfully engrafted, with a 95% confidence interval of 1 out of 211 to 352 *in vitro* expanded ZeMPCs **(Figure 1D)**. However, we note that the ELDA-determined frequency of engrafting cells varies slightly across different ZeMPC derivations, likely due to subtle fluctuations in the efficiency or expansion of these embryo-derived cells in culture. To account for this, we included independent ELDA assessments for both experimental and control conditions in each subsequent transplantation experiment and always used the same freshly derived ZeMPCs for the different treatment groups within an individual experiment. Through these studies, we established a system to perform rapid and cost-effective cell transplant-based screening in zebrafish, which we subsequently deployed to identify chemical compounds that increase the engraftment efficiency of ZeMPCs *in vivo*.

**Figure 1.**
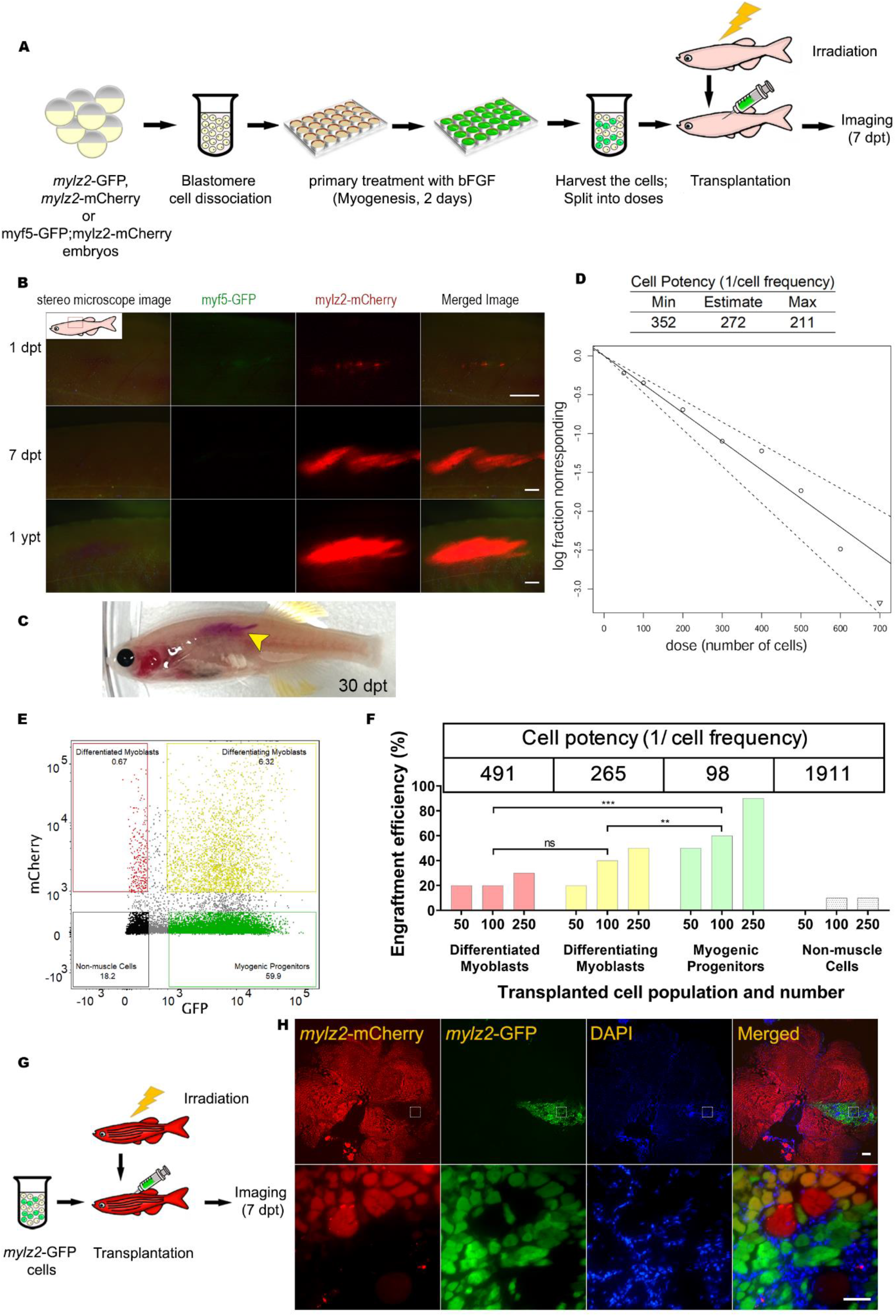
Limiting Dilution Assay (LDA) experimental procedure and statistical analysis. (A) Outline of experimental design. Myogenic progenitors were generated *in vitro* from *mylz2-*GFP or *mylz2*-mCherry embryos. For LDA screening, cells were transplanted into pre-irradiated *casper* recipients (4- to 8-months-old) followed by imaging of the recipient fish at 7dpt. The number of transplanted cells is indicated separately for each experiment. (B) Images of muscle engraftment visualized in the same fish by expression of the muscle-specific (*mylz2-*mCherry) reporter. 10K cells were transplanted into *casper* fish and were stably observed for as long as 1-year post-transplant. myosin light polypeptide chain 2 (Mylz2, red); dpt: days post transplantation; ypt: year post transplantation. Scale bar 500 μm. n = 3. (C) Engrafted muscle-specific (*mylz2-*mCherry) cells are visible to the naked eye. myosin light polypeptide chain 2 (Mylz2, red). (D) Log-fraction plot of the limiting dilution model fitted to the data in **table S1**. The plot represents the fraction of positive responses as a function of the dose of cells delivered in each transplantation. The triangle marks 100% engraftment efficiency with 700-cell transplantation. The line slope is the log-active cell fraction, and the 2 flanking dotted lines give the 95% confidence interval, as listed in the top table. See **table S1** for numbers. (E) *myf5-*GFP; *mylz2-*mCherry double*-*transgenic line cells were sorted into 4 subsets: *myf5*-GFP cells, *mylz2*-mCherry cells, double positive cells, and double negative cells. Each subset was evaluated for engraftment efficiency via limiting-dilution assay. (F) Muscle progenitor cells (*myf5-*GFP) show superior muscle engraftment in recipient fish (n = 10 per cell number dose and 30 per cell population; ***p < 0.001; **p < 0.01; *p < 0.05; ns., not significant; limiting dilution assay). (G) Experimental design of the donor and muscle cell fusion assay. 1 million *mylz2-*GFP ZeMPCs were transplanted into Tg(*mylz2*-mCherry) recipients (4- to 8-months-old). (H) A cross section of a recipient fish at 7 dpt showing fusion of host and donor muscle fibers at the boundary of the engrafted region. The bottom row shows a zoomed image of the box in the top row. myosin light polypeptide chain 2 (Mylz2, red and green) and Nuclei (DAPI, blue). Scale bar 500 μm.

### *myf5-*expressing myogenic progenitors exhibit superior engraftment efficiency

To define the myogenic cell sub-population(s) that supports high efficiency engraftment in zebrafish muscle, *myf5-*GFP*;mylz2-*mCherry double transgenic embryos were generated, dissociated, and cultured with bFGF. The *in vitro* expanded muscle cells at Day 2 were purified into 4 populations of cells by fluorescence-activated cell sorting (FACS): *myf5-*GFP^+^ *mylz2-*mCherry^-^ cells, *myf5-*GFP^-^ *mylz2-*mCherry^+^ cells, *myf5-*GFP^*+*^ *mylz2-*mCherry^+^ cells, and *myf5-*GFP^-^ *mylz2-*mCherry^-^ cells **(Figure 1E)**. Based on their myogenic marker expression, these sorted cells represent myogenic progenitors, terminally differentiated muscle cells, differentiating myoblasts, and other non-muscle cell types, respectively. Cells from each sorted population were transplanted intramuscularly into 10 individual *casper* recipients at each of 3 doses – 50 cells, 100 cells, or 250 cells – to enable limiting dilution analysis. At 7 dpt, fish were anesthetized and prepared for imaging. The engrafting cell frequency was significantly higher for *myf5-*GFP^+^ *mylz2-*mCherry^-^ cells (1 out of 97.7), indicating that myogenic progenitor cells engrafted more readily relative to the other populations evaluated **(Figure 1F and additional SI item**).

### Transplanted embryo-derived zebrafish muscle cells fuse with recipient muscle cells

To determine the manner by which transplanted donor ZeMPCs contribute to muscle regeneration *in vivo*, whether by *de novo* myogenesis or by fusion with endogenous myocytes, 1 million *in vitro* generated muscle cells from *mylz2-*GFP embryos were transplanted into *mylz2-*mCherry adult fish (**Figure 1G**) to generate a large patch of engrafted tissue. At 7 dpt, the recipient fish were euthanized, and the dissected muscle was fixed, sectioned, stained with DAPI, and prepared for imaging to discriminate donor-derived (GFP^+^), host-derived (mCherry^+^) and hybrid (GFP^+^ and mCherry^+^) muscle cells. A cross section of one recipient fish flank shows both *mylz2*-mCherry-tagged recipient muscle cells and *mylz2*-GFP-tagged donor cells (**Figure 1H)**. Although the cells at the center of the engrafted region expressed only GFP, cells at the border were marked with both GFP and mCherry (**Figure 1H, bottom row)**, indicating that the transplanted muscle cells are capable of both *de novo* myogenesis and fusion with recipient muscle. To further test the *in vivo* fusion capacity of transplanted donor cells, we next mixed a limited number of *in vitro* expanded muscle cells from *mylz2*-mCherry embryos with an equal number of cells from *mylz2*-GFP embryos (1:1 ratio; 100 cells per group). The cell mixture was transplanted immediately into *casper* recipients **(Figure S1A)**. At 7 dpt, the recipients were euthanized, and the dissected muscle was fixed, sectioned, stained with DAPI, and prepared for imaging. A cross section of the recipient fish flank shows both *mylz2*-mCherry and *mylz2*-GFP single-color fibers, as well as double-positive (*mylz2*-mCherry and *mylz2*-GFP) hybrid fibers **(Figure S1B)**. In this experiment, single-color fibers were equivalently distributed among GFP-positive (47%) and mCherry-positive (42%) fibers, with 11% of fibers exhibiting both GFP and mCherry, a clear demonstration of the fusogenic ability of transplanted donor cells **(Figure S1C)**. As has been shown previously (Moore et al., 2016), most fibers (∼90%) expressed either GFP or mCherry, and only a small fraction of transplanted cells fused together to make hybrid (GFP^+^ and mCherry^+^) muscle cells.

### Screening for compounds that improve myogenic progenitor cell engraftment efficiency

Lipids have been reported to enhance cell migration and homeostasis in blood and muscle tissue (Cencetti et al., 2014; Lahvic et al., 2018; Li et al., 2015; Oh et al., 2016), and so we elected to screen mainly bioactive lipids in our efforts to identify compounds that enhance ZeMPCs engraftment efficiency *in vivo*. Myogenic progenitor cells were treated for 4 hours with one of the 187 unique lipid compounds contained in the ICCB Known Bioactives Library **(Table S2)**. Compound-treated cells were then washed, harvested and transplanted intramuscularly into both flanks of 5 different *casper* recipient fish, with transplants performed at each of 3 transplanted cell doses (25, 75 or 150 cells). In these studies, recipient fish received split-dose irradiation of 15 Gy at 2 days and 1 day prior to transplantation to suppress any immune responses to the transplanted cells. Engraftment efficiency was measured by imaging *casper* recipients at 7 dpt **(Figure 2A)**. Any compounds that increased engraftment efficiency (ELDA-calculated cell potency) by at least 2 times that of the simultaneously assessed control group (receiving the same concentration of DMSO as the compounds’ vehicle – 0.1% or 1%) were further evaluated using an irradiation-free transplantation model (Moore et al., 2016), which, due to the timing of its generation, was available only for the secondary screening approach. For secondary screening, we transplanted compound-treated cells into *prkdc*^*D3612fs*^ *casper* fish, which are deficient in mature T and B cells, and assayed engraftment efficiency in the recipient fish 7 days post-transplantation. We rank-ordered the compounds according to ELDA-calculated cell potency and picked the top 2, lysophosphatidic acid (LPA) and niflumic acid (NFA), which showed the greatest effect in increasing ZeMPC engraftment efficiency in both assay systems (**Figure 2B**), for further investigation. Notably, both LPA and NFA also increased ZeMPCs engraftment efficiency in competitive transplantation assays, compared to DMSO-treated control cells (**Figure S2**).

**Figure 2.**
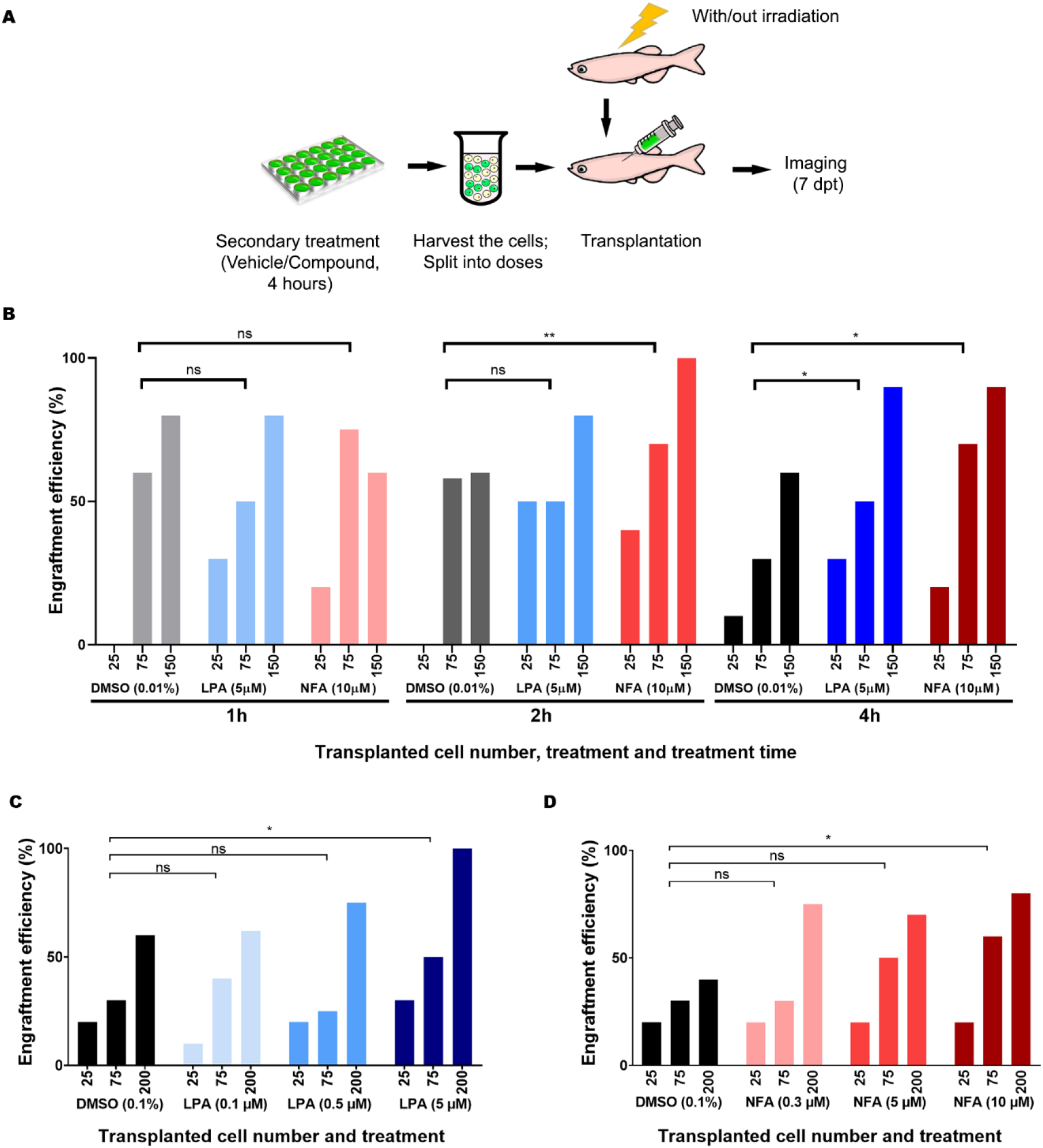
*Ex vivo* exposure to NFA or LPA enhances the engraftment efficiency of zebrafish muscle cells *in vivo*. (A) Outline of experimental strategy. ZeMPCs were incubated with lipids for 4 hours at 28.5°C, followed by washing out of the media and drugs, harvesting the cells, and splitting into 3 cell doses (25, 75 or 200 cells/recipient) for transplantation. Treated cells were transplanted into each side of 5 pre-irradiated *casper* recipient fish or 5 non-irradiated *prkdc*-mutant recipient zebrafish (4- to 8-months-old), followed by imaging the recipient fish at 7dpt. (B) Engraftment efficiency as assessed across different compound exposure times. LPA enhances the engraftment efficiency of ZeMPCs treated for 4 hours, while NFA enhances the engraftment efficiency after 2 and 4 hours of exposure. See also **figure S3** for fold increases and cell potency values. (C and D) Engraftment efficiency of LPA-treated (C) or NFA-treated (D) ZePMCs assessed across different compound concentrations, as indicated. DMSO concentration for all groups including experimental controls and as vehicle for NFA and LPA was 0.1%. (n = 10 per cell number dose and 30 per treatment. **p < 0.01; *p < 0.05; ns., not significant; limiting dilution assay).

### Dose titration of NFA and LPA for enhancing myogenic progenitor cell engraftment efficiency

To further assess the effects of LPA and NFA on ZeMPC engraftment, we treated cells with each compound for different durations and at different concentrations **(Figure 2 an S3)**. Figure 2B compares the engraftment efficiencies under these different conditions; Figure S3A shows the cell potencies and fold increased values and compares engraftment efficiencies of different durations for different treatments. Longer DMSO treatment time resulted in lower engraftment efficiency, where 1 out of 104, 141, and 186 transplanted cells were engrafted after 1, 2, and 4 hours of DMSO treatment, respectively **(Figure S3A)**. These data suggest that under control conditions, ZeMPCs lose engraftment potential during prolonged *in vitro* incubation. In contrast, LPA-treated ZeMPCs showed progressively increased engraftment efficiency, over time and at each experimental time point, when compared to controls. Treatment of ZeMPCs for 1, 2 or 4 hours with LPA increased the engraftment efficiency to 1.11, 1.73 and 2.3 times that of DMSO-treated cells, respectively **(Figure 2B and S3A)**. Similarly, while one-hour NFA-treatment did not alter ZeMPC engraftment, engraftment efficiencies were increased to 2.85 and 2.64 times higher than DMSO-treated cells after 2 or 4 hours of treatment, respectively **(Figure 2B and S3A)**. We also tested engraftment efficiencies of cells treated with DMSO (as vehicle control), or different concentrations of LPA or NFA **(Figure 2C, 2D, S3B and S3C)**. Treatment of ZeMPCs with 5 μM LPA or 10 μM NFA yielded significantly higher (p<0.05) engraftment efficiencies relative to vehicle-treated controls **(Figure 2C and 2D)**—2.61 times higher engraftment efficiency with 5 μM LPA treatment and 2.53 times higher engraftment efficiency with 10 μM NFA treatment **(Figure S3B and S3C)**. Treatment with 0.1 μM LPA or 0.3 μM NFA did not alter the engraftment efficiency of the treated cells (**Figure 2C and 2D**). Furthermore, we found longer time and higher concentration treatment did not result in better engraftment efficiency in comparison to 4 hours treatment with optimal concentrations of LPA or NFA (data not shown). Together, these data confirm the positive effects of both LPA and NFA on engraftment by ZeMPCs and indicate the optimal treatment time (2-4 hours) to drive robust *in vivo* myogenic contributions from transplanted cells treated with these compounds.

### NFA and LPA have additive effects on engraftment efficiency

To determine whether a combination of NFA and LPA might show additive or synergistic effects on ZeMPC engraftment efficiency, we combined different concentrations of NFA and LPA for pre-transplantation treatment. Using the limit dilution assay described above, we found that treatment with 0.1 μM LPA, a concentration equivalent to serum LPA levels in humans (Michalczyk et al., 2017), did not change ZeMPC engraftment efficiency. Engraftment of myogenic progenitors treated with 0.3 μM NFA similarly showed no difference in engraftment rate compared to that of vehicle-treated controls. However, cells treated with either 0.3 μM LPA in combination with 1 μM or 10 μM NFA, or with 0.3 μM NFA in combination with 0.3 μM, 1 μM or 10 μM LPA, showed higher engraftment efficiencies relative to experimental controls (**Figure 3**). Indeed, exposure to 0.3 μM NFA combined with 0.3 μM LPA significantly increased engraftment efficiency compared to vehicle alone, while treatment with the same concentrations of these compounds individually had minimal effects (see **Figure 2C and D**). These data suggest that NFA and LPA have additive effects on myogenic progenitor activity, since combination of NFA and LPA at concentrations that are ineffective individually increases ZeMPC engraftment efficiency.

**Figure 3.**
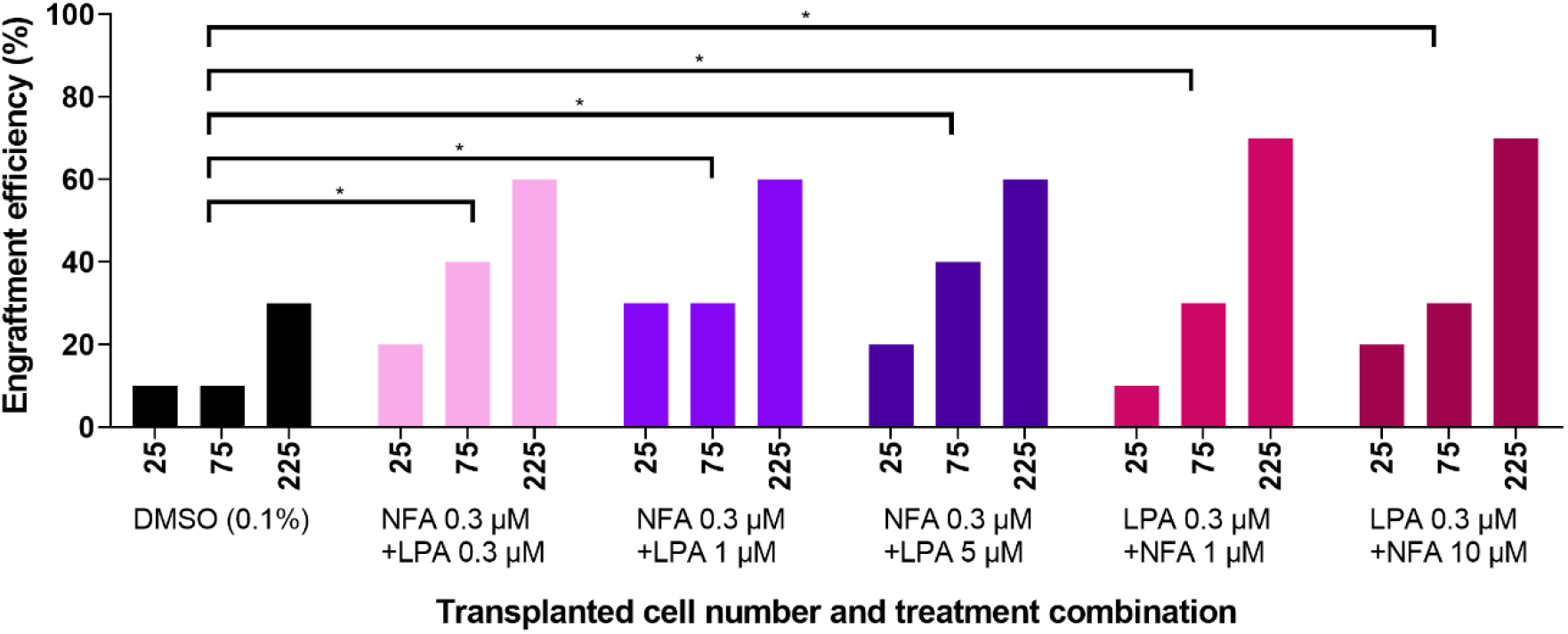
NFA and LPA show an additive effect on muscle progenitor cell engraftment efficiency in zebrafish. Engraftment efficiency in *prkdc*-mutant recipient zebrafish (4- to 8-months-old) of ZeMPCs treated for 4 hours with the indicated concentrations and combinations of NFA and LPA, or with DMSO (vehicle) as control. DMSO concentration for all groups including experimental controls and as vehicle for NFA and LPA was 0.1%. n = 10 fish per cell number dose and 30 per treatment. (*p < 0.05; limiting dilution assay).

### NFA and LPA treatment increase mouse satellite cell engraftment efficiency

To test whether the effects of NFA and LPA on myogenic progenitor cell engraftment potential might be conserved across vertebrate biology, we next assessed the effects of these compounds on mouse satellite cells transplanted into pre-injured recipient tibialis anterior (TA) muscles. Satellite cells were isolated as CD45^-^Sca1^-^Mac1^-^CXCR4^+^β1-integrin^+^ myofiber-associated cells (Castiglioni et al., 2014; Cerletti et al., 2012; Cerletti et al., 2008; Maesner et al., 2016; Sherwood et al., 2004; Sinha et al., 2014; Xu et al., 2013) from FVB-Tg(CAG-eGFP) transgenic mice using our published protocol (Maesner et al., 2016; Sherwood et al., 2004; Xu et al., 2013) (**Figure 4A**). Sorted mouse satellite cells were exposed to NFA, LPA, or vehicle (DMSO) under the optimal dose conditions established using the fish model (see above; 4 h. treatment at 37°C with 10 μM of NFA or 5 μM of LPA), prior to transplant. Compound-treated cells were then recovered and, after washing out the media containing the compounds (NFA, LPA or DMSO), counted and transplanted at 5000 cells per recipient into the pre-injured TA muscles of non-transgenic FVB hosts. Four weeks after transplantation, recipients were euthanized to harvest the injected TA muscles. The muscle tissue was fixed, sectioned, stained with DAPI and WGA, and imaged to quantify the number of GFP^+^ myofibers (Cerletti et al., 2008). Similar to the fish experiments, both NFA and LPA enhanced the engraftment efficiency of mouse satellite cells in these *in vivo* transplantation assays (**Figure 4B, C and 4D**), indicating conserved effects of these pro-myogenic compounds on mammalian muscle precursor cells.

**Figure 4.**
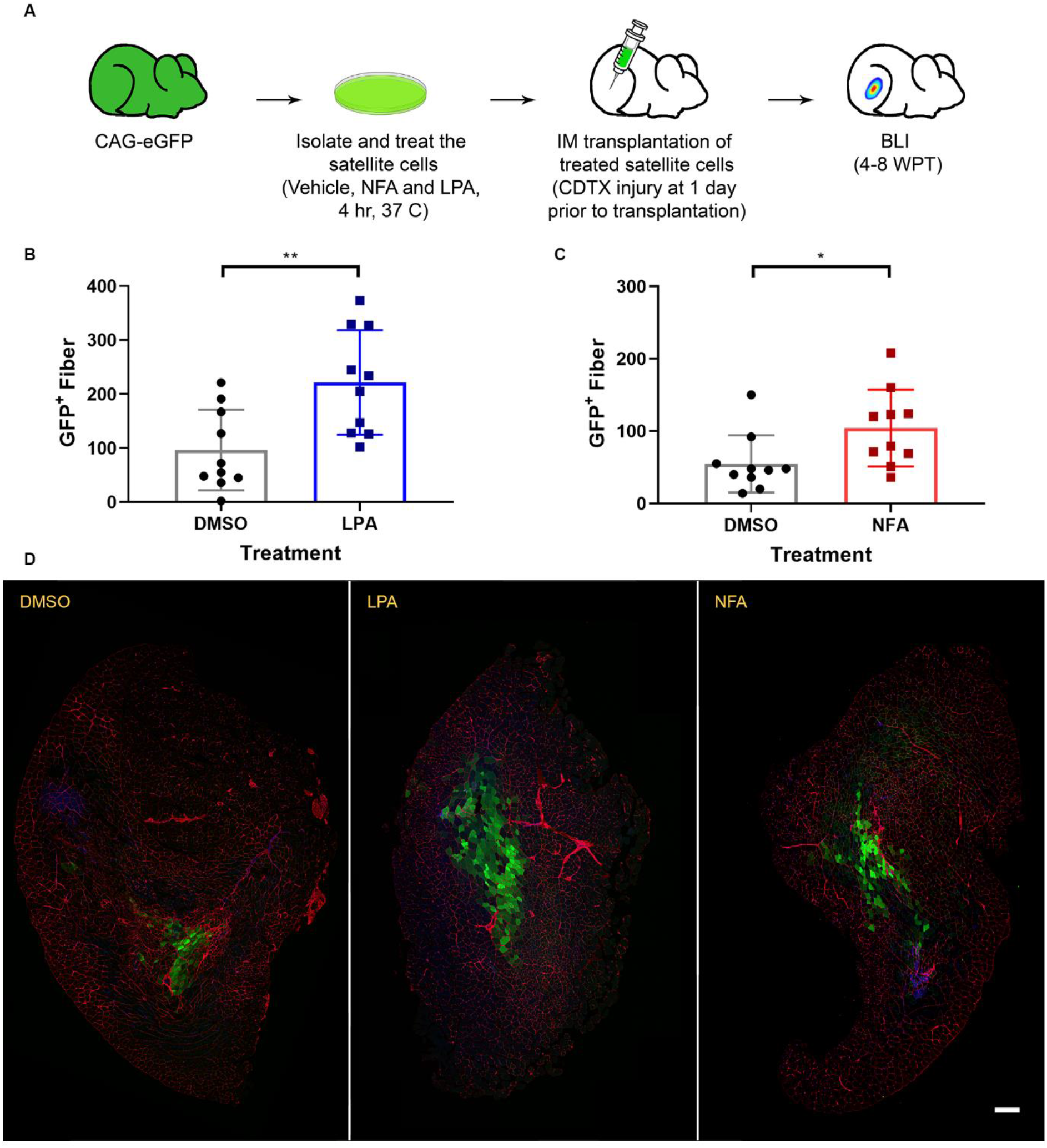
NFA and LPA treatments enhance the engraftment efficiency of mouse satellite cells. (A) Experimental design. (B and C) Engraftment efficiency of 4-hour-treated mouse satellite cells with vehicle, LPA (5 μM) or NFA (10 μM). 5,000 treated cells were injected into preinjured TA muscles and engraftment was measured by counting the number of GFP^+^ myofibers. Data are presented as means ± SEM. Statistical analysis: Two-way ANOVA, and both treatment groups were significantly different from control group (n = 10; **p < 0.01; *p < 0.05). (D) Transverse frozen section of TA muscles injected with 5,000 treated satellite cells. Cell membrane (WGA, red), Nuclei (DAPI, blue), and CAG-eGFP (green). These studies used 8–16-week-old male mice as recipients. DMSO concentration for all groups including experimental control and as vehicle for NFA and LPA was 0.1%. Scale bar 200 μm.

### NFA and LPA regulate expression of muscle development and calcium ion-dependent genes

To gain insight into the mechanisms through which LPA and NFA treatment might enhance donor cell engraftment in vertebrate muscle, we next performed RNA sequencing analysis in treated ZeMPCs and in mouse satellite cells. The sorted zebrafish myogenic progenitors and mouse satellite were exposed to NFA, LPA, or vehicle (DMSO) under the optimal dose conditions (4 h. treatment at 28.5°C for zebrafish myogenic progenitors and 37°C for mouse satellite cells with 10 μM of NFA, 5 μM of LPA or 0.1% DMSO). Compound-treated cells were then recovered and, after washing out the media containing the compounds (NFA, LPA or DMSO), total RNA was isolated for sequencing. Comparison of differential gene expression in NFA-treated or LPA-treated cells to vehicle-treated cells indicated upregulation of calcium-dependent genes in response to these compounds in both ZeMPCs (**Figure S4**) and mouse satellite cells (**Figure 5A and 5B**). Interestingly, treatment of mouse satellite cells with LPA upregulated myoblast fusion related genes, including myomaker (Tmem8c) and Ccl8.

**Figure 5.**
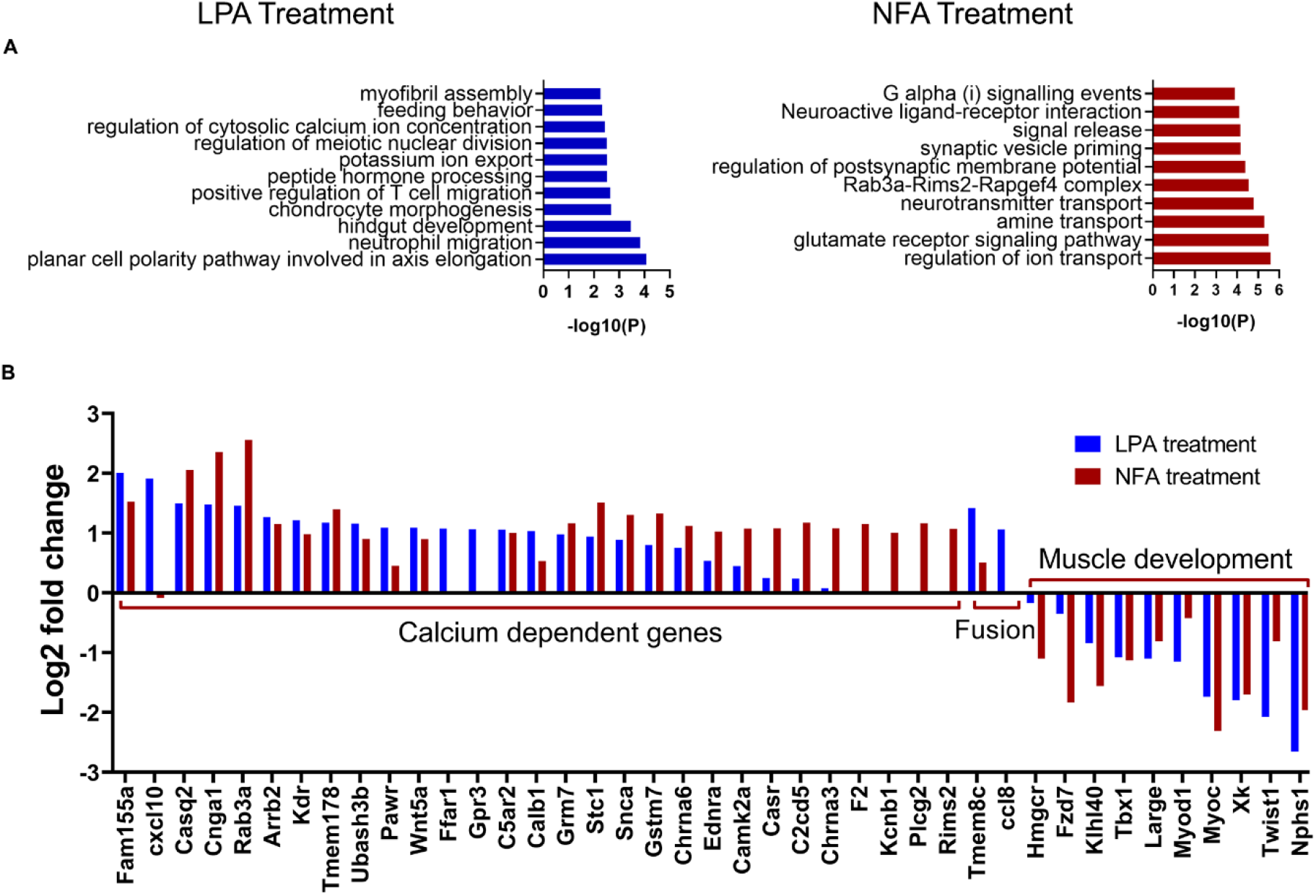
Differential gene expression patterns reveal the upregulation of calcium ion-dependent genes and downregulation of muscle development genes in NFA-treated and LPA-treated mouse satellite cells. **(A)** Gene ontology (GO) enrichment analysis of differentially expressed genes in NFA-treated or LPA-treated mouse satellite cells. **(B)** Log2-fold change in expression of selected genes in LPA and NFA treated mouse muscle stem cells.

To further evaluate the possibility, suggested by these transcriptional studies, that NFA and LPA may act on myogenic progenitors by increasing cytosolic calcium ion concentration, we directly assessed intracellular calcium ion concentration changes in ZeMPCs, in the mouse myogenic cell line C2C12, and in primary mouse satellite cells, in response to added compounds. ZeMPCs and C2C12 cells were incubated with Fura-2, AM, a fluorescent calcium ion indicator, for 45 minutes at 28.5°C and 37°C, respectively, followed by imaging. 20 seconds after imaging initiation NFA or LPA was added to the cells and imaging was continued for a total of 2-3 minutes. Time-lapse calcium ion imaging in ZeMPCs **(Figure S5A)** and mouse myogenic cell line C2C12 **(Figure S5B)** shows both NFA and LPA increase the intracellular concentration calcium ions.

To evaluate the change of intracellular calcium ion concentration across different compound concentrations we isolated mouse satellite cells for calcium imaging analysis. First, to establish optimal conditions, cells were sorted by FACS into 384 well plates at 10,000, 5,000, 1,500, or 500 cells per well, with 8 replicates for each cell density. Subsequent incubation with Fluo4 AM, another fluorescent calcium ion indicator, for 45 minutes at 37°C, showed maximal signal intensity at a density of 1,500 cells per well (**Figure S6)**. Based on these results, we next assessed intracellular calcium ion concentration changes in primary mouse satellite cells plated at 1,500/well and exposed to varying doses of NFA and LPA. Cells were again incubated with Fluo4 AM for 45 minutes at 37°C followed by imaging. Different concentrations of LPA, NFA, or both compounds combined, along with DMSO (vehicle control) and ionomycin (direct calcium ionophore), were added 60 seconds after the beginning of imaging to quantify both baseline calcium ion concentrations and calcium flux in response to the different treatment conditions. We calculated calcium concentrations as area under the curve (AUC) of signal intensity and compared these values across different concentrations of each compound (**Figure S7A, B and C**).

Calcium ion concentrations were increased significantly, relative to vehicle-treated controls, in response to 1-5 μM of LPA and in response to 1-10 μM of NFA (**Figure S7C**), which, notably, overlaps with the concentrations determined to provide optimal muscle cell engraftment in zebrafish and in mice (5 μM LPA and 10 μM NFA, **Figures 2** and **S7D**). The AUC of 0.25 μM NFA combined with 0.25 μM LPA showed a boost in intracellular calcium ion concentration in comparison to the response when each of these drugs was given individually at the same concentrations, correlating with the previously documented additive effect of NFA and LPA on engraftment (**Figure 3**). Interestingly, ionomycin as a positive control for calcium ion influx in mouse satellite cells can be considered as a positive control for muscle cells engraftment, since it also increased the engraftment efficiency of treated ZeMPCs (**Figure S7D**), consistent with a direct, mechanistic role for increased calcium signalling in enhancing the myogenic activity of transplanted muscle progenitors.

### Transplantation of NFA-treated or LPA-treated ZeMPCs improves swimming performance in dystrophic *sapje*-like zebrafish

Finally, to assess the functionality of myogenic precursors exposed to NFA and/or LPA prior to muscle engraftment, we evaluated the swimming performance of *sapje*-like fish before and after transplantation. The *sapje*-like (*sap*^c/100^) fish is a dystrophin mutant and an excellent genetic and phenotypic model of human Duchenne Muscular Dystrophy, demonstrating histological and physiological symptoms of profound muscle degeneration (Guyon et al., 2009). Recipient *sapje*-like fish were irradiated and injected with healthy ZeMPCs, exposed prior to transplant to NFA, LPA or vehicle alone. Cells were injected on both flanks at 3 dorsal points on each side—in the tail, beneath the dorsal fin, and in the trunk—with 25 cells per injection. Immediately prior to transplant, and again 7 days after transplant, we tested muscle function in a modified Blazka-type swim chamber, in which a flow rate of approximately 5.5 L/min was introduced to test swimming performance (**Figure 6**). Recipient fish that were transplanted with NFA-treated or LPA-treated cells, which reproducibly show a higher efficiency of myogenic engraftment (**Figure 2**), performed significantly better in this test when compared to fish engrafted with vehicle-treated cells (**Figure 6B-E**). Because we transplanted only the treated cells, after removal of all compounds, we conclude from these studies that the swimming performance of these transplanted *sapje*-like fish was improved as a result of better engraftment by NFA-treated or LPA-treated ZeMPCs.

**Figure 6.**
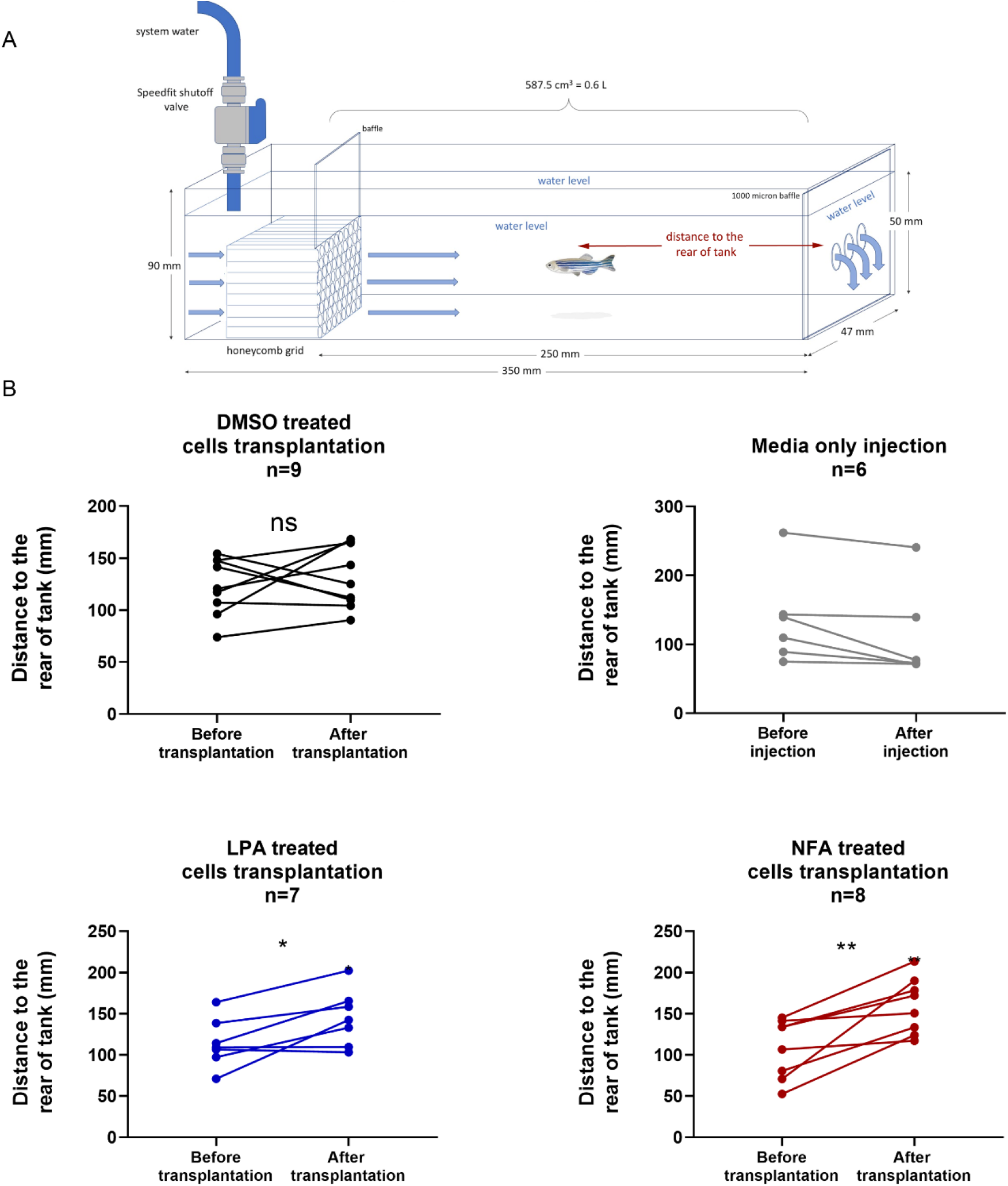
Engrafted cells improve swimming performance in mutant zebrafish. (A) Schematic figure of swimming chamber and analysis strategy. (B) Comparison of *sapje*-like fish (4- to 8-months old) swimming performance before and after transplantation. DMSO concentration for all groups including experimental control and as vehicle for NFA and LPA was 0.1% (* *p* < 0.05; ** *p* < 0.01; ns., not significant; paired *t*-test).

## Discussion

Using the uniquely quantitative, rapid and integrated cross-species screening system described here, we have uncovered new regulators of vertebrate muscle cell engraftment and established a novel platform for discovery of pro-myogenic molecules that act directly on muscle precursor cells and target a stem cell-specific cellular function (*in vivo* engraftment). Multiple studies, including this one, demonstrate the vastly superior capacity of the most primitive subset of muscle precursor cells to contribute productively to muscle repair in transplantation and regeneration assays, particularly in comparison to their differentiated daughters, despite the fact that the two types of cells often show equivalent proliferative capacity in *ex vivo* culture (Cerletti et al., 2012; Cerletti et al., 2008; Gurevich et al., 2016; Montarras et al., 2005; Sacco et al., 2008). Strategies that rely entirely on *ex vivo* screening for numeric increases in cell number (Billin et al., 2016; Nierobisz et al., 2013) after long term (i.e., days, as opposed to only hours in our study) exposure to added chemicals could miss potent stem cell specific compounds whose effects are obscured by the presence of non-stem cells in these cultures. The outcomes of such screens also are highly influenced by the precise culture conditions used (Cosgrove et al., 2014), and might instead favor the identification of compounds that increase cell numbers overall but do not specifically expand or preserve the unique engraftment functions of the muscle stem cell compartment. On the other hand, the use of classical myogenic engraftment assays as a primary outcomes measure in chemical screening approaches has been eschewed by most in the research community due in large part to the prohibitive costs in terms of the time and resources it would require. Most studies to date that have identified single molecules capable of increasing the *ex vivo* yield of satellite cells or enhancing muscle regenerative potential have instead utilized expanded cells and candidate-based approaches, informed in some cases by genetic or transcriptional screening (Bernet et al., 2014; Kuang et al., 2007; Le Grand et al., 2009; Lean et al., 2019; Price et al., 2014; Tierney et al., 2014).

In this study, we sought to overcome this barrier to direct screening on stem cell specific functions in muscle by developing a cross-species approach, first evaluating the myogenic effects of library compounds in a novel zebrafish transplantation system, and then further assessing “hits” in more standard mouse models. A significant advantage of this approach is the increased throughput afforded by use of the zebrafish system (about 14,000 recipient fish were used in this study) and the greater time efficiency of zebrafish myogenesis, which enables specification of myogenic progenitors within 2-3 days and readout of *in vivo* engraftment within a week. In contrast, analogous mouse and human systems require more than 1 month for each step (Xu et al., 2013). Furthermore, extensive data demonstrating the evolutionary conservation of vertebrate myogenesis and our prior work documenting the strong predictive power of the zebrafish system to identify bioactive molecules that similarly regulate mammalian myogenesis (Xu et al., 2013), provided strong rationale for this cross-species approach. We further increased the specificity and rigor of this platform by incorporating highly sensitive limit dilution assays for quantitative assessment of engrafting cell frequencies within populations of compound-exposed cells. We undertook an unbiased assessment of all lipid mediators contained within a focused library of bioactive compounds, and by restricting compound exposure to *ex vivo* pre-transplantation treatment of myogenic progenitor cells, removing compounds by washing prior to transplant, we were able to eliminate any potential toxicity or indirect (bystander) effect of the compounds on the host environment. These features resulted in a robust assay system for studying muscle engraftment.

Using this novel *in vivo* cross-system strategy, we screened 187 compounds in zebrafish and selected the top two - niflumic acid (NFA), an FDA-approved non-steroidal anti-inflammatory drug, and lysophosphatidic acid (LPA), a bioactive phospholipid, for further testing in mice. We documented improved engraftment after pre-transplantation treatment with each of these lipid mediators in both zebrafish and mice, and further identified overlapping cell physiological effects and transcriptional consequences of these two compounds. These observations reinforce the notion that our approach optimally leverages evolutionary similarities between vertebrates to enable more efficient drug identification.

LPA is a prominent member of the lysophospholipid (LP) family, an endogenous class of lipid mediators known to act through sets of specific G-protein-coupled receptors (GPCRs) named LPAR1-LPAR6 (Hecht et al., 1996). LPA may pass through the cell membrane (Stapleton et al., 2011) and may also be generated from membrane phospholipids (Budnik and Mukhopadhyay, 2002). Prior studies have implicated LPA in regulating intracellular calcium ion concentrations, consistent with the results presented here, as well as in the control of cell migration, adhesion, survival, development, and proliferation (Ishii et al., 2004; Sheng et al., 2015; Xu et al., 2008; Ye and Chun, 2010). In studies of cancer cells, LPA’s effects on cell migration have been attributed to its stimulation of Rho activation and actin depolymerization via increases in intracellular calcium ion concentration (Ishii et al., 2004; Kim and Adelstein, 2011). In muscle cells, LPA also has been shown to stimulate the migration and proliferation of cultured myoblasts (Cencetti et al., 2014), with its effects mediated in part through sphingosine kinase and sphingosine-1-phosphate receptors; however, the impact of LPA on the regenerative activities of muscle satellite cells *in vivo* have not previously been assessed. The studies presented here reveal that LPA exposure significantly enhances the engraftment capacities of muscle progenitor cells from both zebrafish and mice, concomitant with increases in cytoplasmic calcium concentration and alterations of myogenic gene expression.

Like LPA, NFA exposure also enhanced the engraftment activities of muscle progenitors across species and induced similar calcium flux and transcriptional alterations. Unlike LPA, which is produced endogenously in fish and mice, NFA is a synthetic compound whose cellular receptor remains unknown. NFA is a member of the non-steroidal anti-inflammatory drug (NSAID) family and can inhibit both phospholipase A2 as well as cyclooxygenases such as COX-2. NFA is frequently used in the treatment of rheumatoid arthritis to reduce pain and suppress inflammation; however, its potential utility for augmenting the contributions of transplanted myogenic progenitors to regenerating muscle has not previously been assessed. Interestingly, prior studies testing the *in vivo* effects of NSAID administration on endogenous muscle repair have reported both pro-regenerative (Oh et al., 2016) and anti-regenerative (Ho et al., 2017) impact. This suggests differences in the specific actions of individual NSAIDs and/or pleiotropy of their action on different cell types *in vivo*. In this regard, the *ex vivo* exposure and transplantation strategy implemented here presents a distinct advantage, avoiding such complications by restricting compound exposure to only myogenic precursor cells and thereby revealing the particular effects of these molecules on stem cell-specific regenerative activities.

In our studies, we found that both NFA and LPA trigger rapid increases in intracellular calcium ion concentration within treated muscle progenitors, consistent with prior observations in other cell types (Liantonio et al., 2007; Xu et al., 2008). Notably, these effects of LPA and NFA were additive, potentially reflecting previously reported differences in the mechanisms by which they impact calcium flux. In particular, while NFA triggers increases in intracellular calcium by inducing the release of calcium ions from intracellular stores, increases of intracellular calcium seen with LPA treatment have been attributed to influx of calcium ions from extracellular sources (Poronnik et al., 1992; Rao et al., 2003). We posit that exposure to NFA and LPA together enables mobilization of calcium from both intracellular and extracellular depots, resulting in an additive effect.

We also found that LPA increases the expression of myoblast fusion regulators and effectors, including Ccl8 (Griffin et al., 2010) and Myomaker (also known as Mymk and Tmem8c) (Millay et al., 2013; Quinn et al., 2017). Such effects may promote the increased contribution of LPA-treated cells to the formation of multinucleated fibers in transplant recipients. Moreover, in addition to its effects promoting myogenic cell fusion, Ccl8 also has been implicated in promoting myogenic differentiation (Ge et al, 2013) and as a positive regulator of the release of sequestered calcium ions into the cytosol (Richardson et al., 2000). Thus, induction of Ccl8 by LPA may further enhance the calcium mobilizing and pro-myogenic activities of this compound.

As an ultimate test of the ability of compounds discovered in our screening system to stimulate productive contributions to regenerative myogenesis by transplanted muscle precursor cells, we examined the impact on dystrophic *sapje*-like fish of engraftment with NFA-treated or LPA-treated cells, simulating a clinical cell therapy scenario. Engraftment with either NFA-treated or LPA-treated muscle progenitors resulted in significant improvements in swimming performance of the recipient fish, in comparison to control fish receiving equivalent doses (25 cells per injection at each of 6 different injection sites) of vehicle-treated ZeMPCs. Given our consistent observation that LPA and NFA treatment increases the engraftment efficiency of donor ZeMPCs, together with the fact that compound exposure was restricted to the pre-transplantation period, with careful washing to remove any residual compound prior to cell injection, we conclude from these studies that the improvement in swimming performance reflects an improved engraftment efficiency, with no direct effect on the recipient muscle tissue. Such *ex vivo* chemical treatment approaches could be applied in clinical cell therapy, as a strategy to boost the per cell regenerative output of *ex vivo* expanded or pluripotent cell derived human muscle progenitors.

In summary, this work establishes a novel screening platform for the discovery of pro-myogenic compounds that act specifically on muscle stem cells and target a muscle stem cell-specific function – myogenic engraftment after *in vivo* transplantation. Applying this platform to interrogate a relatively understudied class of bioactive molecules, lipid mediators, we discovered two – NFA and LPA – that enhanced the *in vivo* engraftment efficiency of both *in vitro* expanded embryo-derived zebrafish muscle cells and freshly isolated mouse muscle satellite cells. We further showed that LPA and NFA also significantly increase intracellular calcium ion concentration in zebrafish muscle cells, C2C12 mouse muscle cells, and mouse satellite cells. We discovered that the combination of these two lipids at low concentrations had an additive effect and enhanced muscle progenitor cell engraftment efficiency. Finally, we evaluated the functionality of transplanted cells and showed that transplantation of NFA- and LPA-treated ZeMPCs improved zebrafish muscle function as measured by swimming performance, in comparison to vehicle-treated cells. This study indicates that pre-transplantation drug treatment can be used to improve muscle cell therapy approaches and rescue muscle function through progenitor cell engraftment in muscular dystrophies.

## Experimental Procedure

### Fish housing, husbandry and Fish Transgenic Lines

All fish used in the experiment were housed in 3.5L tanks with recirculating water and kept at 10 fish per liter. Water quality was kept at a constant 1250 μS, pH 7.5, and a water temperature of 28.5°C. Fish were fed Gemma Micro 500 at approximately 5% body weight per day and housed under a photoperiod of 14hr-light, 10hr-dark cycle. All fish were bred and housed following standard zebrafish husbandry(Westerfield, 2007). In this study, we used the following transgenic fish lines: Tg(*myf5*-GFP), Tg(*mylz2*-GFP), and Tg(*mylz2-*mCherry) for zebrafish blastomere culture and ZeMPCs generation. Tg(MHC-*casper*) and Tg(*prkdc*^*D3612fs*^) (*casper*-strain) as recipients in the screen experiment and Tg*(sapje*-like (*sap*^c/100^) for function study.

### Culture and myogenesis of dissociated zebrafish blastomere cells

Myogenic progenitor cells were generated *in vitro* from *mylz2-*GFP, *mylz2-*mCherry (RRID:ZFIN_ZDB-ALT-071016-1) or *myf5-*GFP;*mylz2-*mCherry embryos using our published protocols (Xu et al., 2013) with minor changes. 20-30 embryos were used to dissociate the blastomere cells and grown in a zESC medium composed of 70% LDF medium (50% Leibowitz’s L-15 (Gibco), 35% DMEM (Gibco), and 15% Ham’s F-12 (Gibco)), with 20% embryo extract and 10% FBS; these were supplemented with 2 ng/ml recombinant human FGF basic protein (Sigma-Aldrich), 15 mM sodium bicarbonate, 15 mM HEPES (Gibco),1% L-glutamine (Gibco), 10 nM sodium selenite (Sigma-Aldrich), 1% N2 (Gibco), 2% B27 (Gibco), and 0.1 mg/ml Primocin (Invivogen). Cells were cultured at 28°C without CO_2_ for 48 hours.

### Chemical treatments and zebrafish *in vivo* screen

Lipids from the ICCB Known Bioactives Library (enzolifesciences.com/BML-2840/iccb-known-bioactives-library) (see **Table S2**) were diluted at a 1:100 ratio in transplantation media (zESC medium without bFGF and FBS) and added to prewashed *in vitro* expanded muscle cells. NFA (Cayman and Santa Cruz Biotechnology) and LPA (Cayman and Santa Cruz Biotechnology) were used for secondary screening experiments. DMSO concentration was 1% for the primary screen (in consistent with 1:100 ratio dilution of the library), and 0.1% for the other experiments including secondary screen, competitive transplantation, mouse experiments and functional study. Zebrafish cells were incubated with lipids for 4 hours at 28.5°C, followed by washing out the media and drugs, harvesting the cells, and splitting into 3 cell doses for transplantation. 4- to 8-month-old *casper* recipients received split-dose irradiation of 15 Gy each, either 2 days or 1 day before transplantation. For LDA screening, cells from each group were split into 3 doses and transplanted into each side of 5 pre-irradiated *casper* recipient fish or 5 non-irradiated *prkdc*-mutant recipient zebrafish. The *prkdc*-mutant zebrafish line was available only after we initiated the primary screen. Because irradiation could interfere with muscle cell engraftment (Doreste et al., 2020), we used the *prkdc*-mutant zebrafish for repeat experiments and to confirm of the validity of the primary screening system. At 7 dpt, recipients were anaesthetized using a previously described method (Dang et al., 2016) and imaged using a fluorescent stereo microscope (Leica M165 FC). Successful engraftment was defined as the presence of *mylz2*-GFP^+^/ *mylz2*-mCherry^+^ fibers. Variation in different experiments is caused by inherent variability in the *in vitro* ZeMPC derivation procedure, which results in differences in engrafting cell frequency across different derivation attempts. To mitigate the impact of such experimental variation on our screening results, we only compared LDA results for cells derived from the same cultures and included comparison to a vehicle-treated experimental control in every transplantation experiment. Chemical treated cell potency and vehicle-treated cell potency were measured and analysed using the Extreme Limiting Dilution Analysis (ELDA) software (Hu and Smyth, 2009). The ratio of compound-treated cell potency and vehicle-treated cell potency determine the engraftment efficiency fold increase value.

### Zebrafish tissue sampling, staining, and imaging

Adult zebrafish recipients were euthanized at 7 dpt. The dissected body trunks were fixed in 4% paraformaldehyde, followed by cryoprotection with 30% sucrose solution at 4°C overnight. Tissue specimens were embedded with Leica OTC tissue-freezing medium (Leica 14020108926) and rapidly frozen in liquid nitrogen, then sectioned with a cryostat at −25°C (Leica CM1860). The sections were transferred to a room temperature Opaque-coated slide (VWR® Superfrost® Plus Micro Slide 48311-703), followed by air-drying overnight. Sections were demembranated, blocked, and stained with 0.5% Triton X-100, 3% BSA, and DAPI, respectively. Sections were protected by embedding in mounting medium (Vectashield H-1400) and covered with a coverslip. The mounted slides were stored at 4°C in a light-protected condition to preserve the fluorescence before imaging. Images were acquired using Zeiss Axio Scan.Z1 and Zeiss 880 in the Harvard Center for Biological Imaging. The captured images were processed quantitatively with Zen, ImageJ, and MATLAB.

### Functional assay setup and swimming performance measurement

Juvenile zebrafish from a *sapje*-like transgenic line were fin-clipped and genotyped to identify heterozygotes. A flow-through modified Blazka-type swim chamber was created using a horizontal acrylic tank with dimensions of 350mm l x 47mm w x 90mm d. System water, equal in quality to recirculating water in housing tanks, was introduced into one side of the swim chamber at a flow rate of approximately 5.5L min^-1^. Water flow funnelled through a honeycombed grid composed of 50 vinyl tubes (6.35mm 0D, 3.97mm ID x 50mm l). This created a laminar flow through the remaining 250mm length of the swim chamber x 50mm deep x 47mm wide, limiting fish to a swimming area of approximately 587.5 cubic cm = 0.6L. Another 1000-micron baffle was positioned downstream of the tunnel to prevent any fish from flowing out of the three drainage points located at the far end of the swim chamber. Adult *sapje*-like fish were netted out of their holding tanks and placed inside the already flowing swim chamber before and 7 days after transplantation. A high-definition Nikon D3100 digital camera was used to record individual fish’s swimming performance, filmed at 30 frames per second for a total of 3 minutes. If a fish reached a point of fatigue where it could no longer maintain its position in the swim chamber, the fish was swept downstream onto the 1000-micron baffle located at the far end of the tank. A plastic transfer pipette was used to assist the fish off the baffle if it could not free itself and continue swimming. The raw digital file was then analysed for movement within the swim area. The distance from the flow source was quantified using MATLAB (MathWorks; RRID:SCR_001622).

### Mouse Husbandry and handling

FVB-Tg(CAG-eGFP) mice (Stock 003516; RRID:IMSR_JAX:003516), and FVB/NJ mice (Stock 001800) were obtained from the Jackson Laboratory. All mice were housed in the Animal Facility of Harvard University and all the experiments and protocols were performed in compliance with the institutional guidelines of Harvard University. These studies used adult (8–16 weeks of age) male mice.

### Mouse satellite cell isolation, treatment, and transplantation

Mouse satellite cells were isolated as previously described (Maesner et al., 2016; Sherwood et al., 2004; Xu et al., 2013). 25 μl (0.03 mg/ml) of cardiotoxin (CDTX) (Latoxan) was injected 24h prior to transplantation into the TA muscle of recipient mice (FVB/NJ), to stimulate a regenerative response (Cerletti et al., 2008). GFP tagged satellite cells were isolated and plated on a collagen/laminin-coated plate and treated with NFA (10 μM), LPA (5 μM) or DMSO, as the vehicle control, in culture media for 4 hours at 37°C. The culture media contained 78% F10 (GIBCO), 20% horse serum (Atlanta Biologics), 1% penicillin-streptomycin (Gibco), 1% GlutaMAX (Gibco), and 5 ng/ml bFGF (Sigma). After 4 hours, cells were washed with DPBS, harvested, and counted, followed by transplantation of 5000 satellite cells intramuscularly into the CDTX pre-injured TA of FVB recipients.

### Mouse TA sampling, cryosectioning, staining, and imaging

Transplanted TAs were dissected and fixed in 4% paraformaldehyde, followed by cryoprotection with a 30% sucrose solution. Next, the TAs were washed with DPBS and frozen in pre-incubated isopentane in liquid nitrogen for 30 seconds followed by a 30-second incubation in liquid nitrogen. Frozen TA muscles were sectioned using a cryostat at −25°C (Leica CM1850) at 10 μm thickness. Sections were stained with WGA (Thermo Fisher) and DAPI (Thermo Fisher) and embedded in mounting medium (Vectashield, vector laboratories). Images were acquired using Axio Scan.Z1 (Zeiss) and Zeiss 880 in the Harvard Center for Biological Imaging. Captured images were processed quantitatively with ZEN and ImageJ.

### Culture of the mouse myogenic cell line C2C12

Cells were grown at 37°C in growth media: DMEM (Gibco), %10 FBS and 1% penicillin-streptomycin (Gibco).

### Calcium imaging in ZeMPCs and C2C12 cells

ZeMPCs or C2C12 cells were incubated with Fura-2, AM (Thermo Fisher Sci F1225), a cell permeant fluorescent calcium ion indicator, for 45 minutes at 28.5°C, followed by imaging with Celldiscoverer7 Microscope (Zeiss). ZeMPCs were incubated in transplantation media (zESC medium without bFGF and FBS to simulate the transplantation experiment) and C2C12 cells at 37°C in growth media. 20 seconds after imaging initiation the indicated compound(s) or vehicle was added to the cells and imaging was continued for a total of 2-3 minutes.

### Calcium imaging in mouse satellite cells

Satellite cells were isolated from wild-type mice (FVB/NJ) and sorted into 384 well plates in culture media. Cells were incubated overnight at 37°C. The next day, cells were incubated with Fluo-4 AM (Thermo Fisher Sci F14201), a fluorescent calcium ion indicator, for 45 minutes at 37°C, followed by imaging of the entire plate with the FDSS 7000ex functional drug screening system (Hamamatsu) in the BCH assay development and screening facility.

### RNA sequencing

Isolated mouse muscle satellite cells and ZeMPCs were used to prepare RNA after *in vitro* chemical treatment. Total RNA was isolated from 2000 cells per replicate and 3 replicates per treatment by the micro RNeasy kit (QIAGEN) for both mouse muscle satellite cells and ZeMPCs. cDNA was prepared using SMART Seq v4 Ultra Low RNA-Seq kit for 48 reactions (Takara) and a Nextera kit was used for library construction. FASTQ files for samples were processed in tophat-cufflinks workflow in a Linux server operating system to output gene-level abundance estimates and statistical inference as gene-level raw counts. Those raw counts for samples were input into the cuffdiff for differential gene expression analysis. The assigned GEO accession number for the RNA sequencing data is GSE143801.

### Quantification and statistical analysis

All results are presented as raw data or mean ± standard error of the mean (SEM) where indicated. Statistical analysis was performed using the LDA test to measure the number of engrafting cells (potent cells), two-way ANOVA to test the murine engraftment efficiency in different groups and paired t tests to test the swimming performance in the function study (Graphpad Prism®; version 8.3). Sample size, replicate number, treatment concentration and/or treatment duration for each experiment are indicated in the figure legends. Results with p values of less than 0.05 were considered statistically significant: * *p* < 0.05; ** *p* < 0.01; ns., not significant.

## ACKNOWLEDGMENTS

We thank T. M. Schlaeger (Boston Children’s Hospital) for providing the ICCB Known Bioactive Library; D. M. Langenau (Massachusetts General Hospital) for Tg(*myf5*-GFP), Tg(*mylz2*-GFP), Tg(*mylz2-*mCherry), and Tg(*prkdcD3612fs)* (*casper*-strain) line; J. LaVecchio, S. Ionescu, and N. Kheradmand (HSCRB/HSCI Flow Cytometry Core) for flow cytometry support; D. Richardson, Erin Diel, and Christian Hellriegel (Harvard Center for Biological Imaging) for imaging support; L. M. Kunkel (Boston Children’s Hospital) for Tg*(sapje*-like (*sap*^c/100^) line; L. Barrett (Boston Children’s Hospital, the Assay Development and Drug Screening Facility (ADSF)) for calcium imaging support; and The Bauer Core Facility at Harvard University for high throughput RNA sequencing. This work was supported by NIH grants U01 HL100402 (to A.J.W.), an HHMI Early Career Award (to A.J.W), a grant from the Paul F. Glenn Foundation for Medical Research (to A.J.W) and R24 grant from the NIH (to L.I.Z). This work was conducted with financial contributions from Harvard University and its affiliated academic healthcare centers. The content is solely the responsibility of the authors and does not necessarily represent the official views of Harvard University, its affiliated centers, or the National Institutes of Health.

## Author Contributions

S.T., A.J.W. and L.I.Z. designed the experiments, interpreted the data, and wrote the paper. S.T., A.R., E.G., S.A.K., V.S.C., M.E.M., K.A.M. and I.A. conducted the experiments and analysed the results. I.A. designed the swimming chamber. H.F. provided the computer code. S.Y. performed RNA-seq analysis. All authors discussed the results and commented on the manuscript.

## Supplementary Data

**Supplementary table 1.**
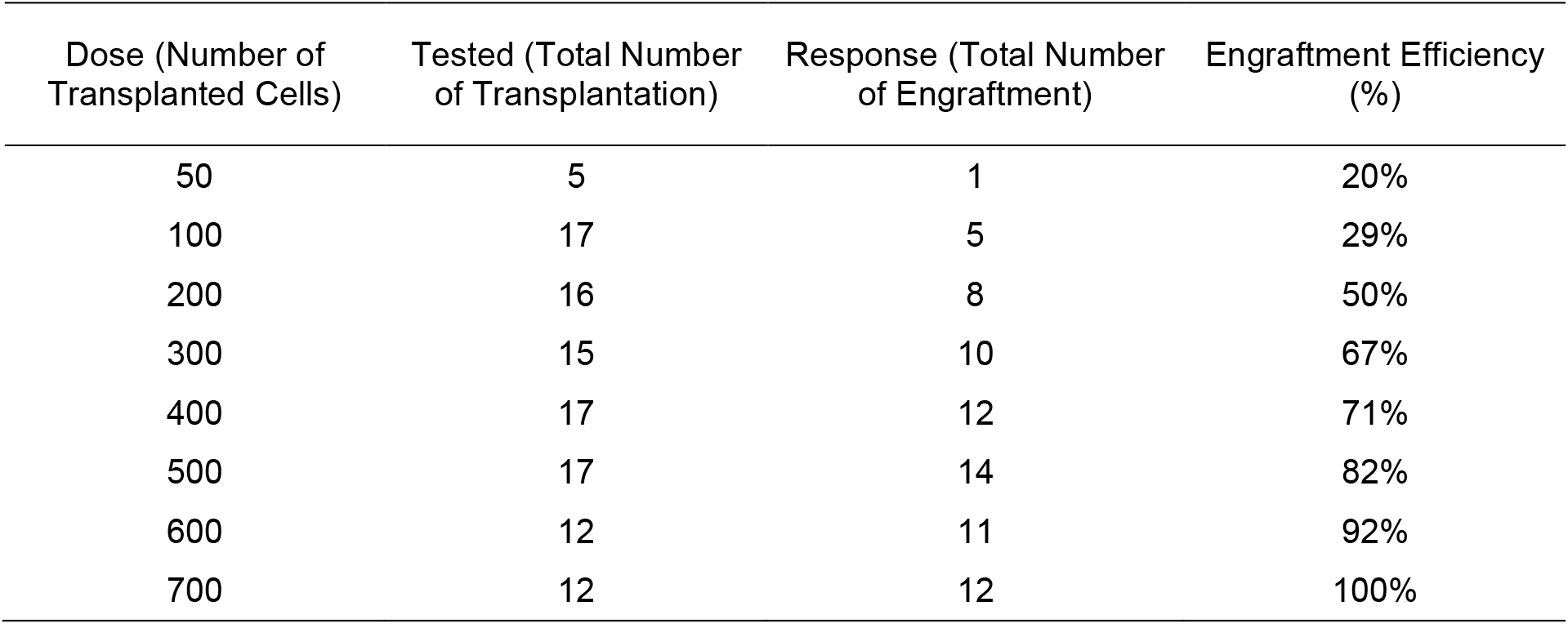
Limiting dilution data showing the frequency of transplanted ZeMPCs, tested fish, responses, and calculated engraftment efficiency for ELDA.

**Supplementary table 2. ICCB known bioactive library.** The table lists detailed information of the 187 lipid compounds included in the primary screen, including name, formula, concentration in the library, molecular weight, etc.

**Figure S1.**
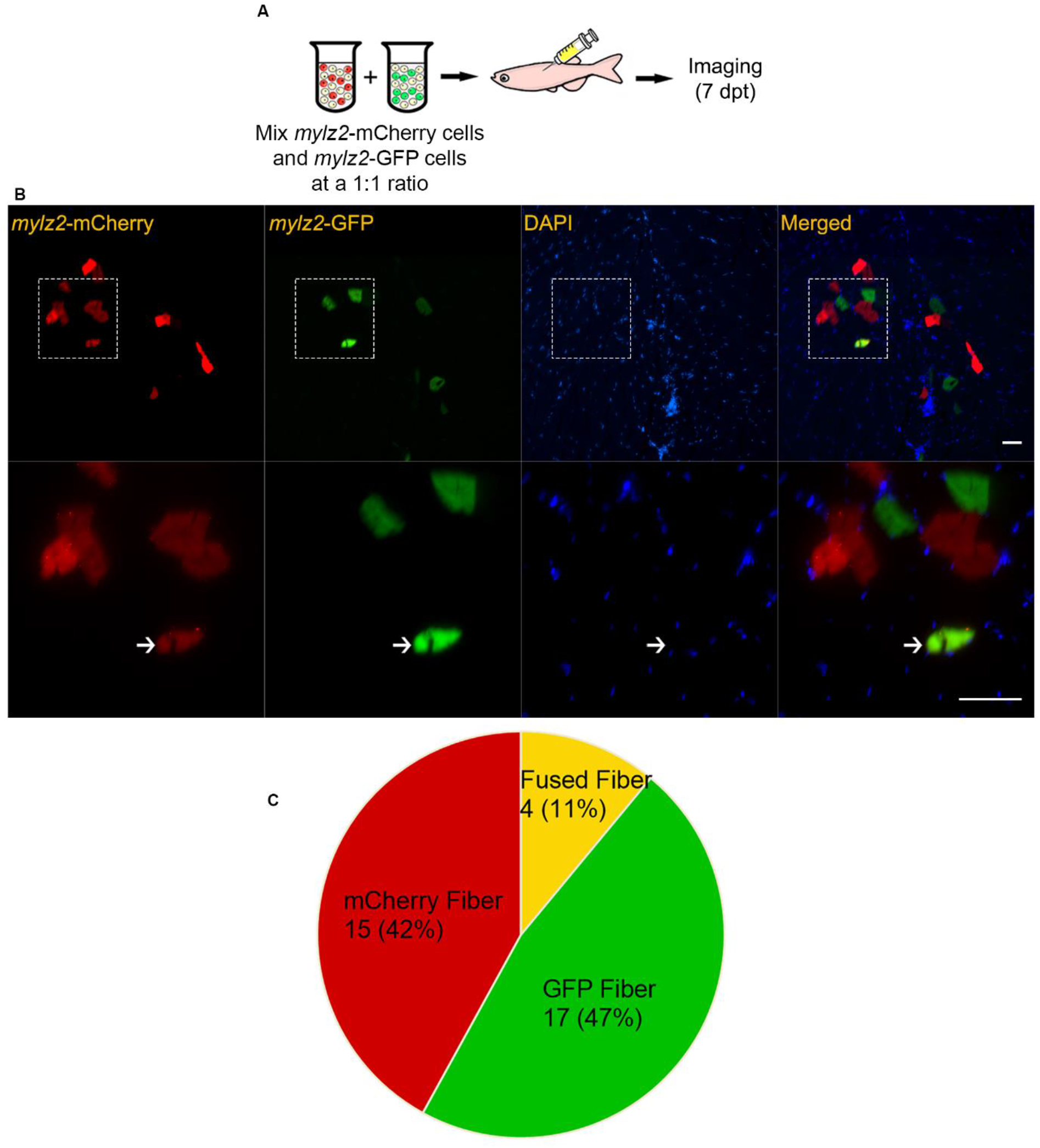
Evaluation of *in vivo* fusion capacity of ZeMPCs. (A) Experimental design. 100 *mylz2-*GFP ZeMPCs and 100 *mylz2-*mCherry ZeMPCs were co-transplanted into *prkdc*-mutant recipient zebrafish. (B) Cross section of a *prkdc*-mutant recipient zebrafish (4- to 8-months old) at 7dpt, demonstrating fusion of the two genotypes of donor cells (hybrid fibers: GFP^+^mCherry^+^). The bottom row shows a zoomed image of the box in the top row. Myosin light polypeptide chain 2 (Mylz2, red and green), Nuclei (DAPI, blue). (C) Quantification of *in vivo* fusion of donor muscle cells in the recipient fish (n = 5 fish). Scale bar 200 μm.

**Figure S2.**
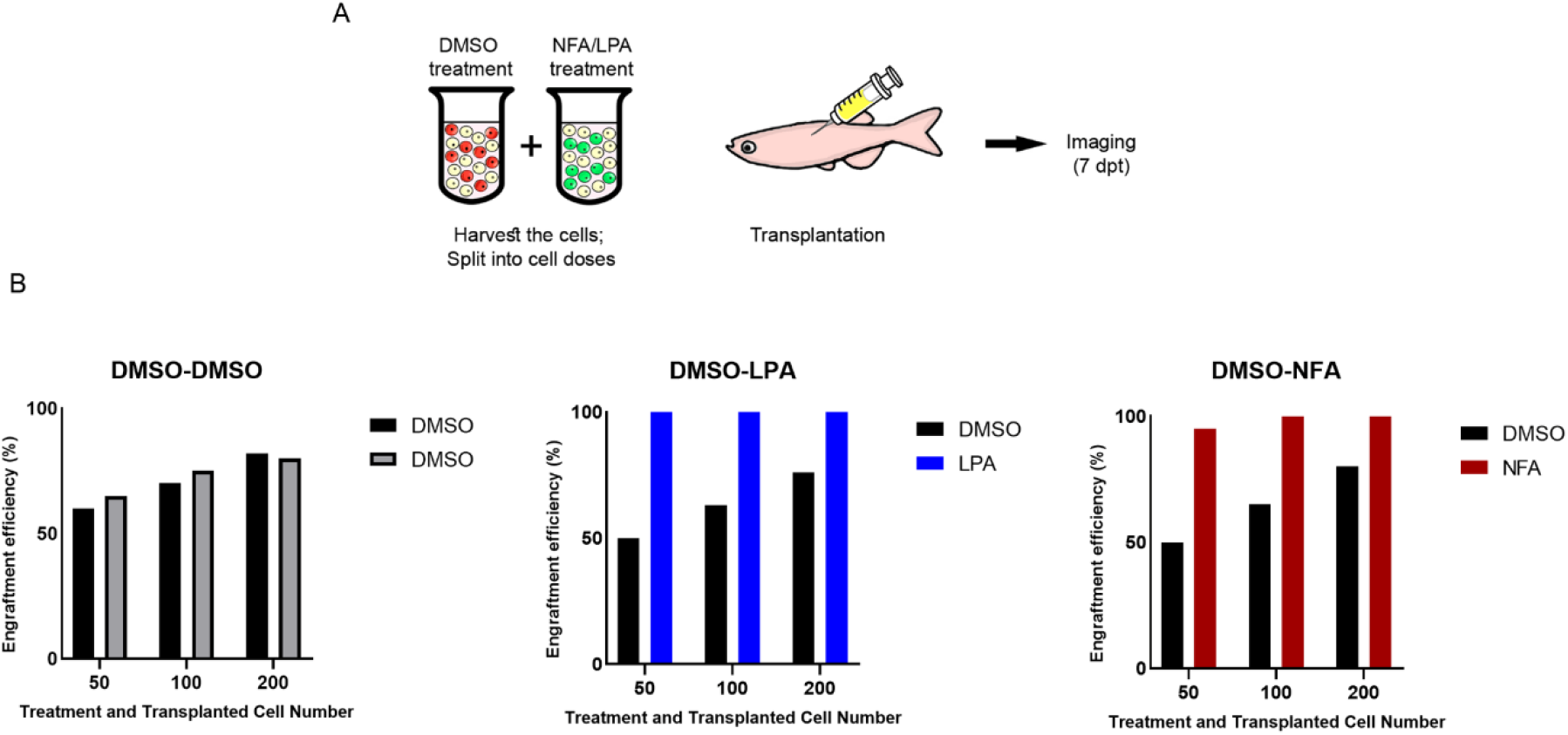
ZeMPC competitive transplantation assays using different combinations of chemical-treated or DMSO-treated ZeMPCs. (A) Experimental design. ZeMPCs from *mylz2*-GFP and *mylz2*-mCherry donors were co-injected into 4- to 8-months-old *prkdc*-mutant recipient zebrafish and imaged at 7 dpt to assess the engraftment of *mylz2*-mCherry (pre-transplantation treatment with DMSO) and *mylz2*-GFP (pre-transplantation treatment with DMSO, LPA or NFA) cells. (B) Engraftment efficiency of equal numbers of DMSO-treated mCherry^+^ cells vs. DMSO-treated GFP^+^ cells (left), DMSO-treated mCherry^+^ cells vs. LPA-treated GFP^+^ cells (middle) and DMSO-treated mCherry^+^ cells vs. NFA-treated GFP^+^ cells (right). DMSO-treated *mylz2*-mCherry ZeMPCs and *mylz2*-GFP ZeMPCs showed similar engraftment (grey bars), whereas pre-treatment with LPA (blue bars) or NFA (red bars) provided a competitive advantage (n = 10 per cell number dose and 30 per treatment).

**Figure S3.**
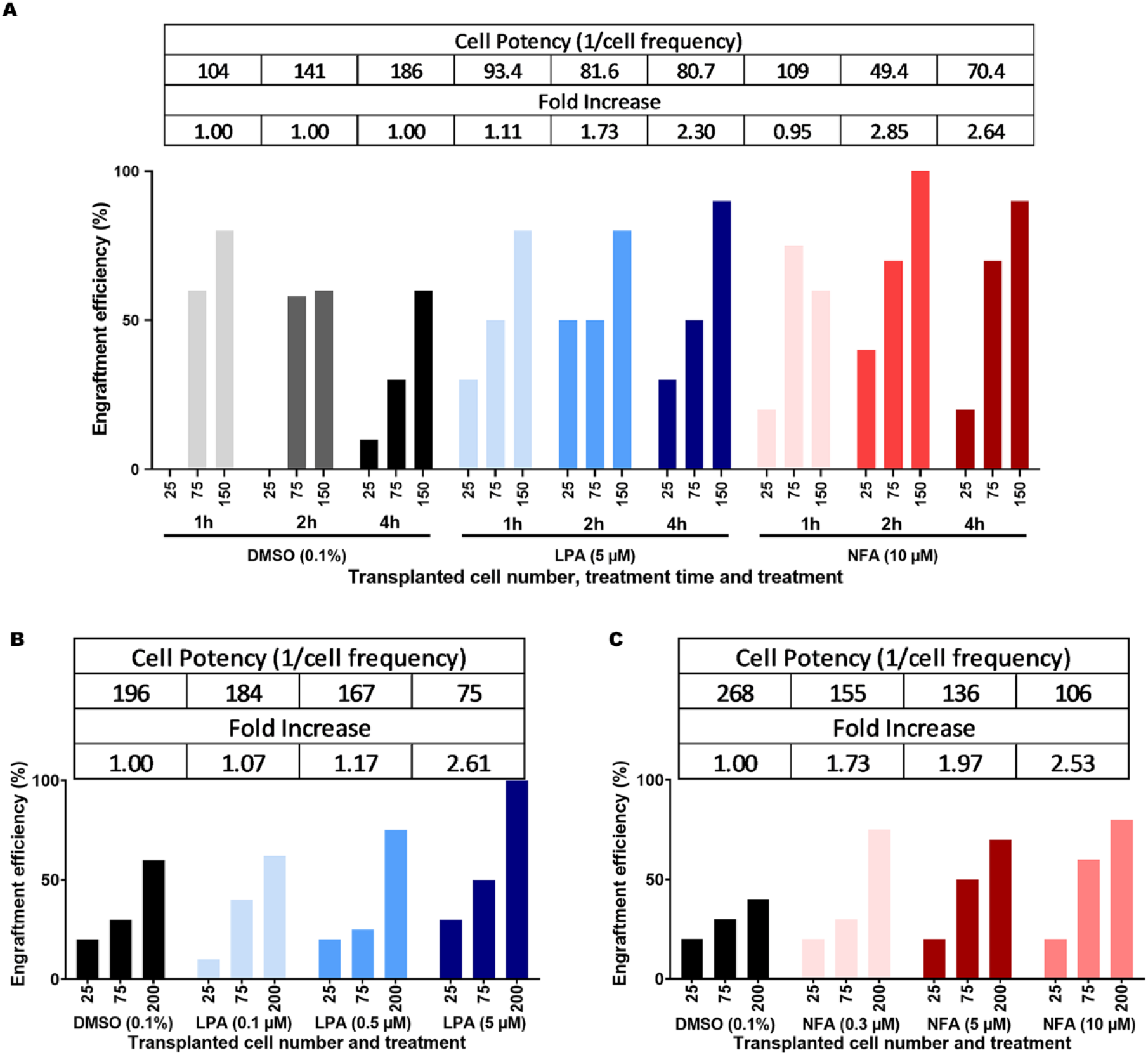
NFA and LPA treatments enhance the engraftment efficiency of zebrafish muscle cells *in vivo*. The purpose of the LDA is to test whether different treatments produce the same engrafting cell proportions. Using ELDA software, the treated cell potency, which indicates the number of transplanted cells required for engraftment (top table), was measured. Fold increase of the engraftment efficiency was calculated as the ratio of the LPA- or NFA-treated cell potency and DMSO-treated cell potency; fold increase of the engraftment efficiency describes how much the engraftment efficiency changed in comparison to the experimental control group (DMSO) for 1, 2, and 4 hours of LPA/NFA treatment in comparison to 1, 2, and 4 hours of DMSO treatment, respectively (bottom table) (n = 10 per cell number dose and 30 per treatment)

**Figure S4.**
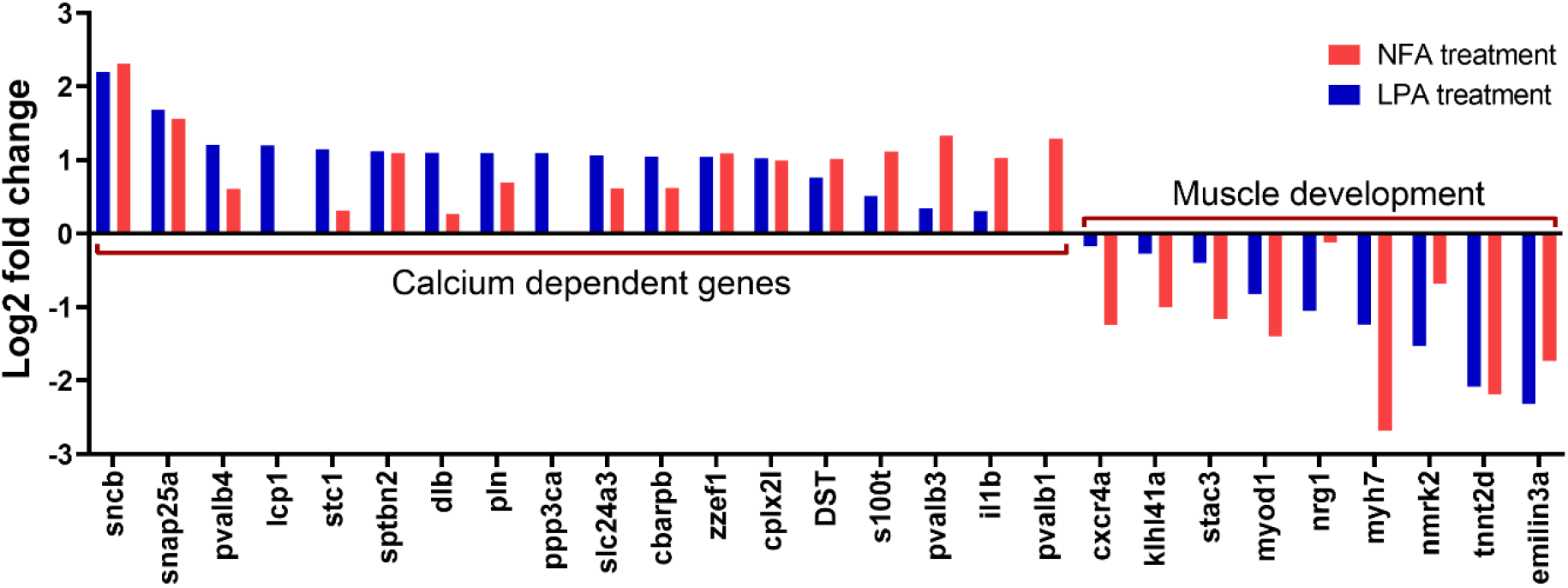
Differential gene expression pattern reveals upregulation of calcium ion-dependent genes in treated ZeMPCs. Log2-fold change in expression of selected genes in LPA and NFA treated ZeMPCs.

**Figure S5.**
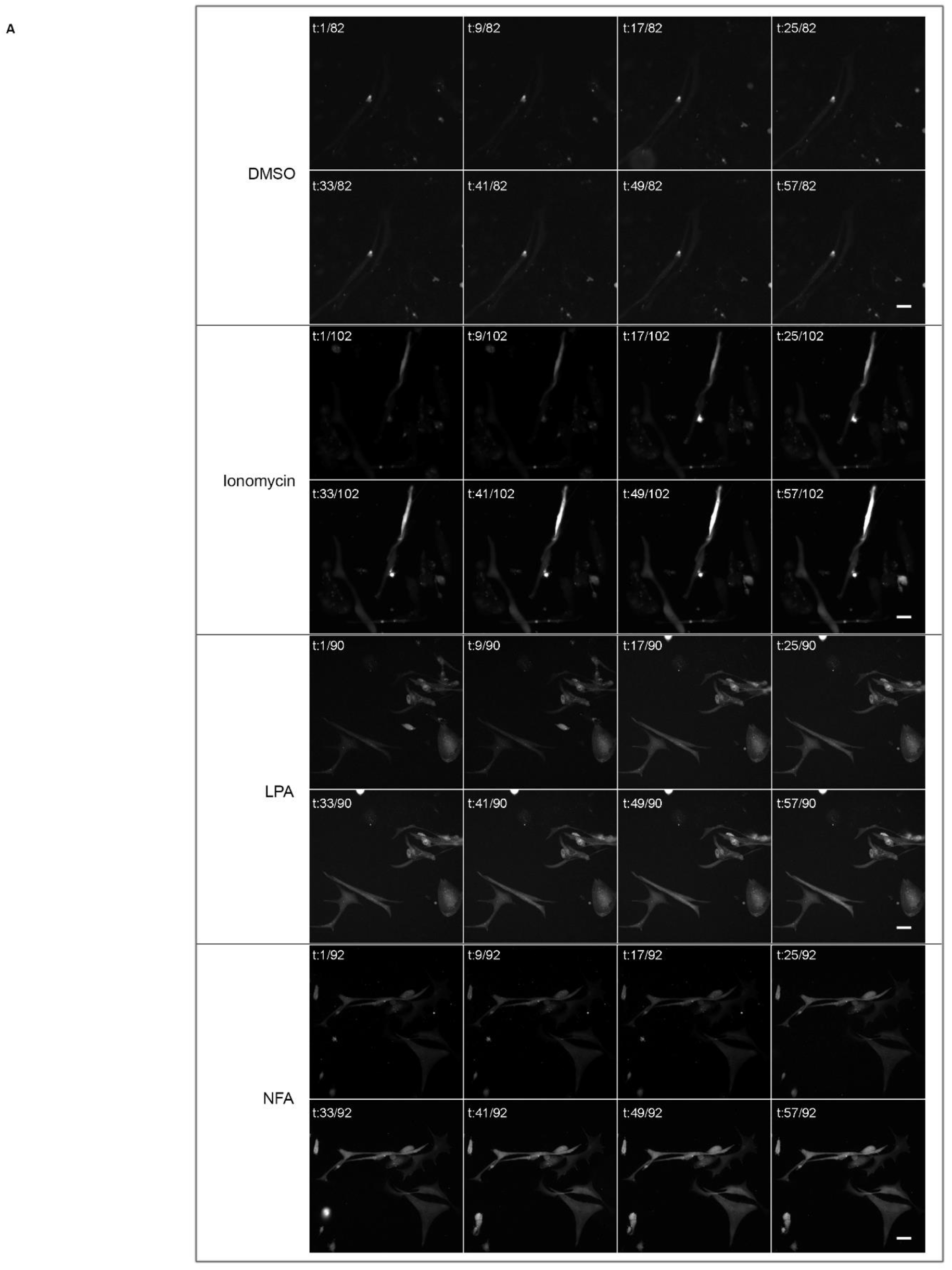
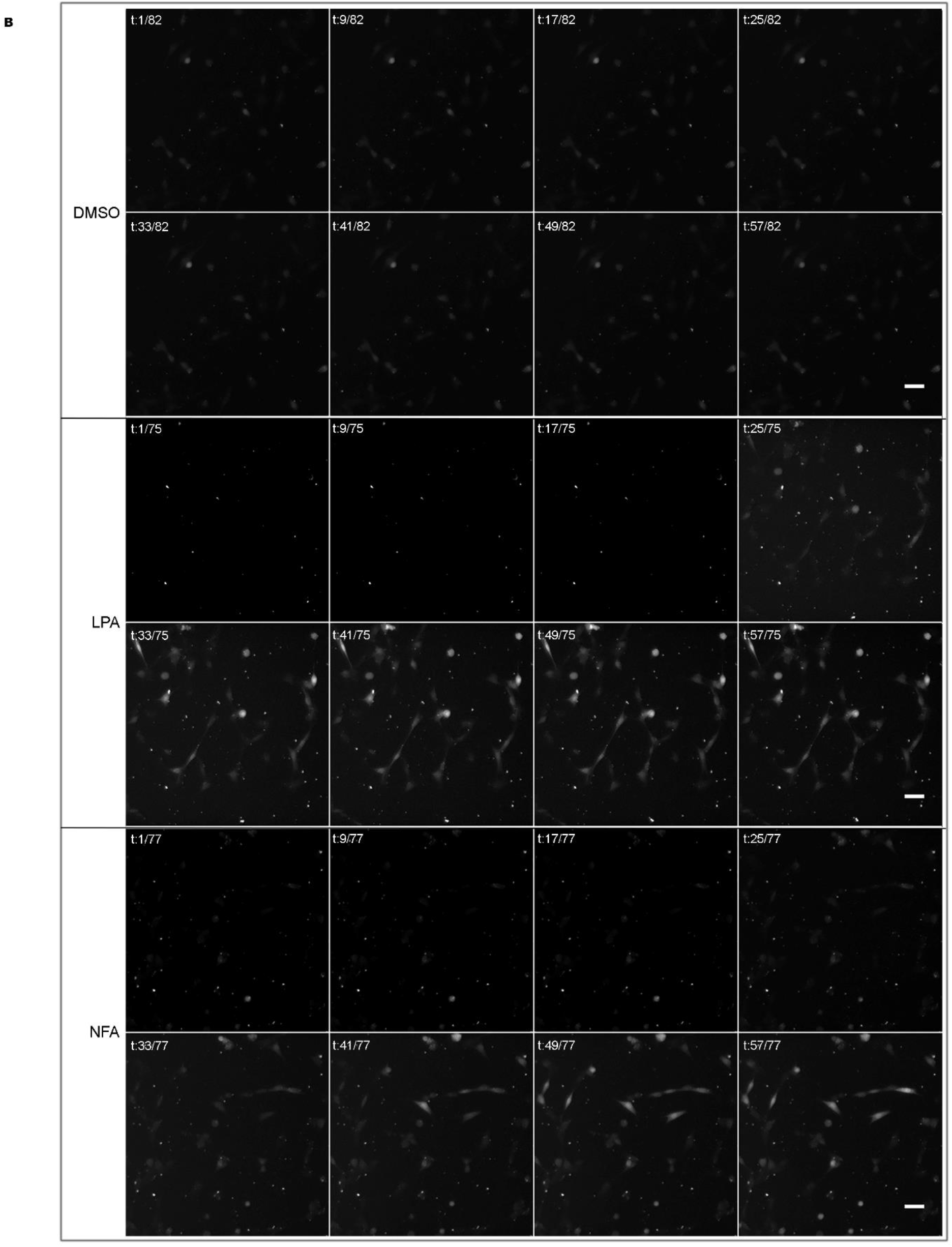
Intracellular calcium ion concentration increases in response to NFA and LPA in *in vitro* expanded zebrafish muscle cells and the mouse myogenic cell line C2C12. Fura-2, AM treated (A) ZeMPCs were exposed to 10 μM NFA, 5 μM LPA, 10 μM Ionomycin (as a positive control) and DMSO (as vehicle control) and (B) C2C12 cells were exposed to 10 μM NFA, 5 μM LPA and DMSO (as vehicle control) to visualize effects on intracellular calcium ion concentration using time-lapse imaging. Comparing signal intensity before loading the treatment (t: 1 and 9 for ZeMPCs and t: 1, 9 and 17 for C2C12 cells) to signal intensity after loading the treatment, reveals NFA, LPA and ionomycin increase the intracellular calcium ion concentration. Scale bar 50 μm.

**Figure S6.**
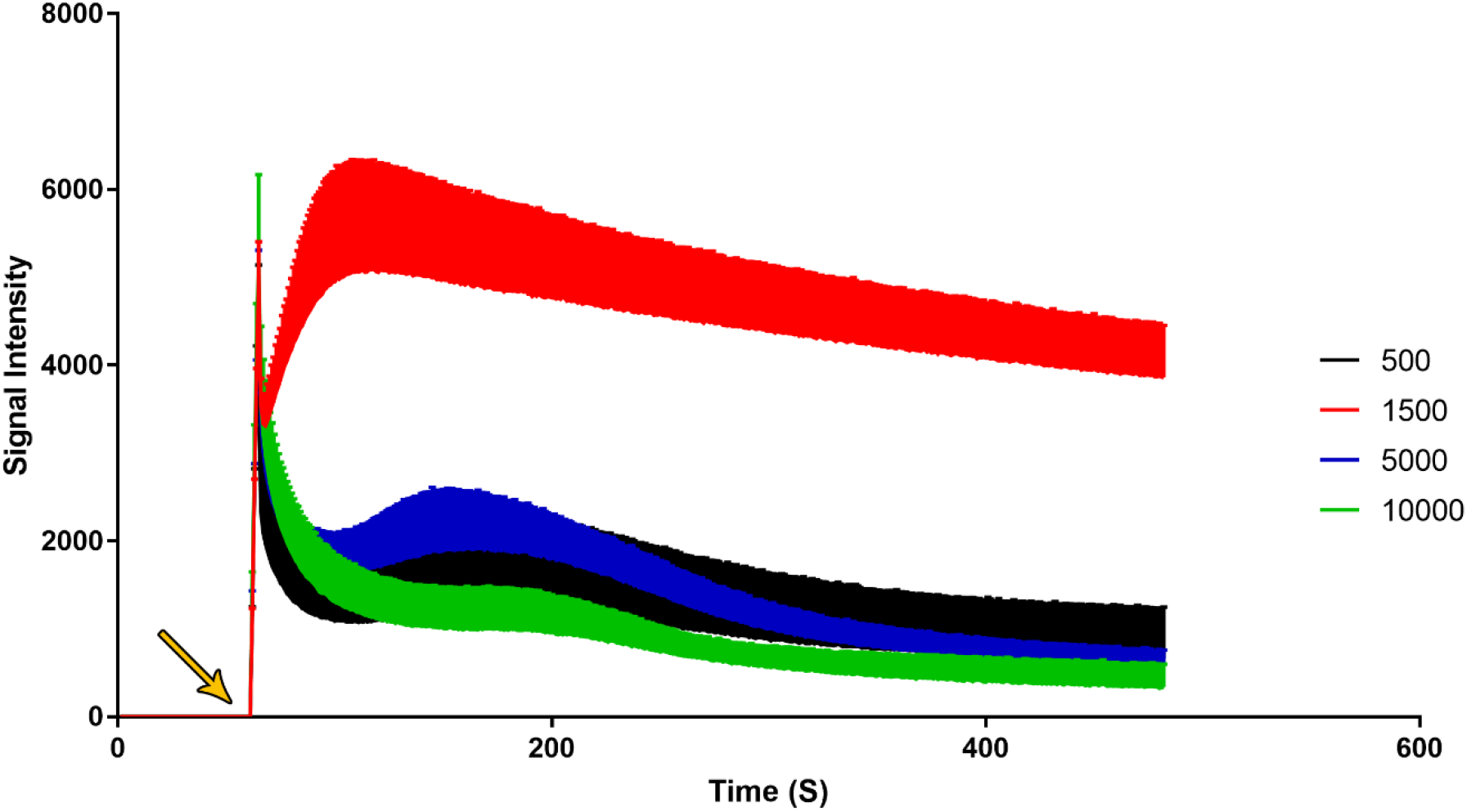
Satellite cell density optimization for calcium imaging. Signal intensity was measured for different initial cell densities: 10,000 (green), 5,000 (blue), 1,500 (red), and 500 (black) mouse satellite cells, which were seeded in 384 well plate (n = 8). The graph reports the signal intensity of the 8 replicates for each cell density (Mean ± SD). Media containing free calcium ions was added to all wells at the timepoint marked with an arrow. The sorted satellite cells in 384 well plates were incubated with Fluo4 AM for 45 minutes at 37°C, followed by imaging.

**Figure S7.**
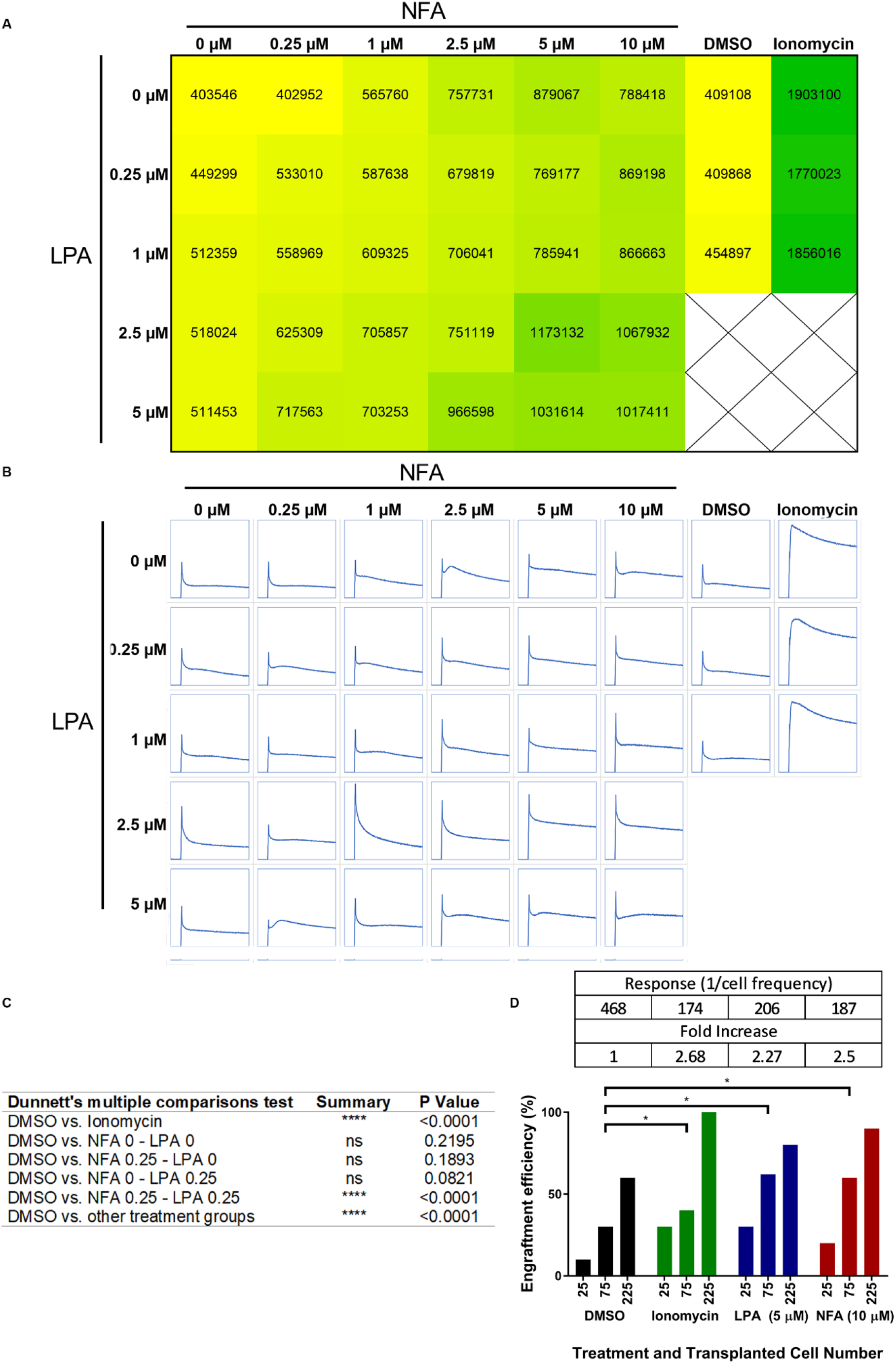
LPA and NFA increase intracellular calcium ion concentrations in mouse satellite cells. (A and B) Measurement of calcium ion transients in Fluo4 AM-loaded mouse satellite cells. (A) Each box reports the area under curve (AUC) inside the well containing the indicated concentrations of NFA, LPA, DMSO, and/or Ionomycin. (B) Each box reports signal intensity (y-axis)-time curve (x-axis) inside the well containing the indicated concentrations of NFA, LPA, DMSO, and/or Ionomycin. (C) Statistical analysis for intracellular calcium ion concentration; Similar to LDA experiment the treatment with 0.25 μM LPA or NFA did not alter the intracellular calcium ion concentration, while the combination treatment with 0.25 μM LPA and 0.25 μM NFA enhanced the intracellular calcium ion concentration (P < 0.0001; Two-way ANOVA and multiple comparisons test). (D) Ionomycin (green bars), a positive control for increased intracellular calcium ion concentrations, enhances the engraftment efficiency of muscle cells in 4- to 8-month-old *prkdc*-mutant zebrafish relative to vehicle-treated controls (black bars), similar to LPA (blue bars) and NFA (red bars). n = 10 fish per cell number dose and 30 per treatment (*p < 0.05; limiting dilution assay).

## Additional SI item

Extreme limiting dilution assay (ELDA) is a technique to measure the number of engrafting cells (potent cells) in a cell population, which perform the desired effect. In this work, we aimed to test the effect of pre-transplantation treatment with specific compounds on muscle engraftment (desired effect) of the donor cells (potent cells). Using LDA, we calculated the number of potent cells for each treatment and tested if the treatment changed the number of potent cells significantly. Below, we show the limiting dilution data, lower, estimate and upper cell potency calculation, pairwise statistical analysis, and log-fraction plot of the limiting dilution model for **figure 1F**.

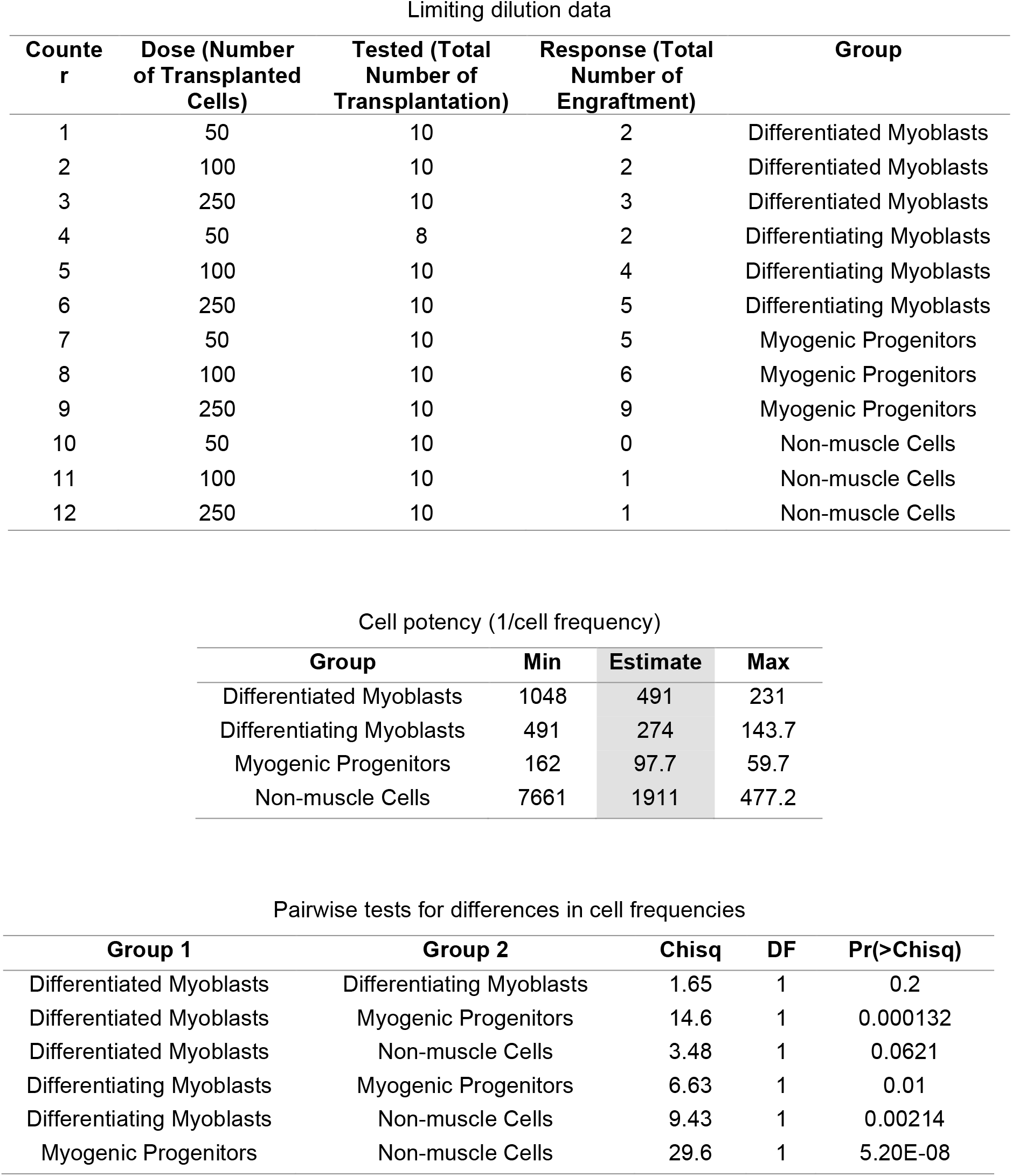

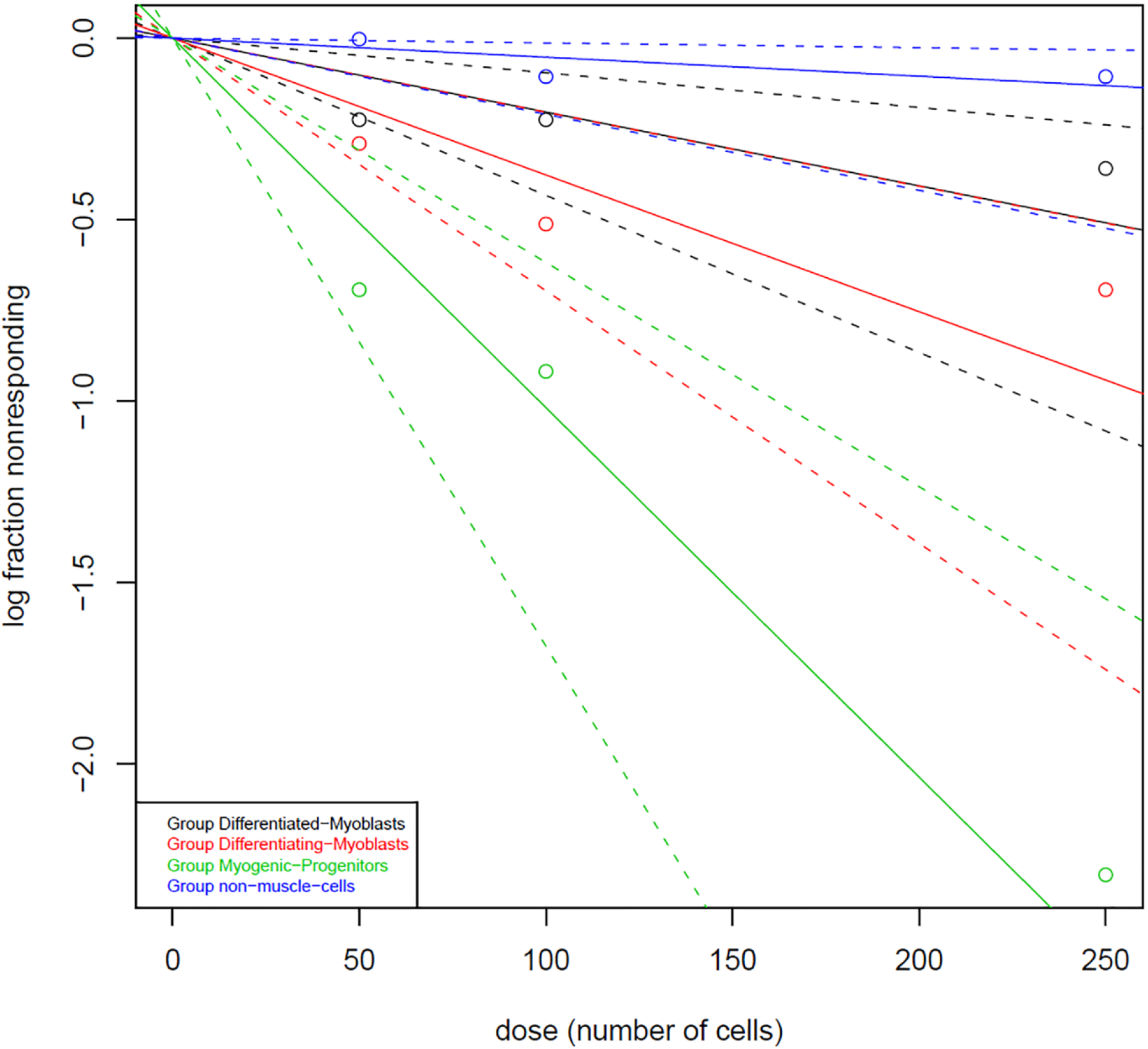

## References

Bernet, J.D., Doles, J.D., Hall, J.K., Kelly Tanaka, K., Carter, T.A., and Olwin, B.B. (2014). p38 MAPK signaling underlies a cell-autonomous loss of stem cell self-renewal in skeletal muscle of aged mice. Nature medicine 20, 265–271.

Berridge, M.J., Lipp, P., and Bootman, M.D. (2000). The versatility and universality of calcium signalling. Nature Reviews Molecular Cell Biology 1, 11.

Billin, A.N., Bantscheff, M., Drewes, G., Ghidelli-Disse, S., Holt, J.A., Kramer, H.F., McDougal, A.J., Smalley, T.L., Wells, C.I., Zuercher, W.J., et al. (2016). Discovery of Novel Small Molecules that Activate Satellite Cell Proliferation and Enhance Repair of Damaged Muscle. ACS Chemical Biology 11, 518–529.

Bischoff, R. (1975). Regeneration of single skeletal muscle fibers in vitro. The Anatomical record 182, 215–235.

Budnik, L.T., and Mukhopadhyay, A.K. (2002). Lysophosphatidic Acid and Its Role in Reproduction. Biology of Reproduction 66, 859–865.

Carlson, B.M. (1973). The regeneration of skeletal muscle. A review. The American journal of anatomy 137, 119–149.

Carlson, B.M., and Faulkner, J.A. (1996). The regeneration of noninnervated muscle grafts and marcaine-treated muscles in young and old rats. The journals of gerontology Series A, Biological sciences and medical sciences 51, B43–49.

Castiglioni, A., Hettmer, S., Lynes, M.D., Rao, T.N., Tchessalova, D., Sinha, I., Lee, B.T., Tseng, Y.H., and Wagers, A.J. (2014). Isolation of progenitors that exhibit myogenic/osteogenic bipotency in vitro by fluorescence-activated cell sorting from human fetal muscle. Stem cell reports 2, 92–106.

Cencetti, F., Bruno, G., Blescia, S., Bernacchioni, C., Bruni, P., and Donati, C. (2014). Lysophosphatidic acid stimulates cell migration of satellite cells. A role for the sphingosine kinase/sphingosine 1-phosphate axis. FEBS Journal 281, 4467–4478.

Cerletti, M., Jang, Y.C., Finley, L.W., Haigis, M.C., and Wagers, A.J. (2012). Short-term calorie restriction enhances skeletal muscle stem cell function. Cell stem cell 10, 515–519.

Cerletti, M., Jurga, S., Witczak, C.A., Hirshman, M.F., Shadrach, J.L., Goodyear, L.J., and Wagers, A.J. (2008). Highly Efficient, Functional Engraftment of Skeletal Muscle Stem Cells in Dystrophic Muscles. Cell 134, 37–47.

Collins, C.A., Olsen, I., Zammit, P.S., Heslop, L., Petrie, A., Partridge, T.A., and Morgan, J.E. (2005). Stem cell function, self-renewal, and behavioral heterogeneity of cells from the adult muscle satellite cell niche. Cell 122, 289–301.

Conboy, I.M., Conboy, M.J., Smythe, G.M., and Rando, T.A. (2003). Notch-mediated restoration of regenerative potential to aged muscle. Science (New York, NY) 302, 1575–1577.

Cosgrove, B.D., Gilbert, P.M., Porpiglia, E., Mourkioti, F., Lee, S.P., Corbel, S.Y., Llewellyn, M.E., Delp, S.L., and Blau, H.M. (2014). Rejuvenation of the muscle stem cell population restores strength to injured aged muscles. Nature medicine 20, 255–264.

Cox, G.A., Cole, N.M., Matsumura, K., Phelps, S.F., Hauschka, S.D., Campbell, K.P., Faulkner, J.A., and Chamberlain, J.S. (1993). Overexpression of dystrophin in transgenic mdx mice eliminates dystrophic symptoms without toxicity. Nature 364, 725–729.

Dang, M., Henderson, R.E., Garraway, L.A., and Zon, L.I. (2016). Long-term drug administration in the adult zebrafish using oral gavage for cancer preclinical studies. Disease models & mechanisms 9, 811–820.

Doreste, B., Torelli, S., and Morgan, J. (2020). Irradiation dependent inflammatory response may enhance satellite cell engraftment. Scientific Reports 10, 11119.

Ervasti, J.M. (2007). Dystrophin, its interactions with other proteins, and implications for muscular dystrophy. Biochimica et Biophysica Acta (BBA) - Molecular Basis of Disease 1772, 108–117.

Fukada, S.-i., Higuchi, S., Segawa, M., Koda, K.-i., Yamamoto, Y., Tsujikawa, K., Kohama, Y., Uezumi, A., Imamura, M., Miyagoe-Suzuki, Y., et al. (2004). Purification and cell-surface marker characterization of quiescent satellite cells from murine skeletal muscle by a novel monoclonal antibody. Experimental Cell Research 296, 245–255.

Griffin, C.A., Apponi, L.H., Long, K.K., and Pavlath, G.K. (2010). Chemokine expression and control of muscle cell migration during myogenesis. J Cell Sci 123, 3052–3060.

Gurevich, D.B., Nguyen, P.D., Siegel, A.L., Ehrlich, O.V., Sonntag, C., Phan, J.M., Berger, S., Ratnayake, D., Hersey, L., Berger, J., et al. (2016). Asymmetric division of clonal muscle stem cells coordinates muscle regeneration in vivo. Science (New York, NY) 353, aad9969.

Guyon, J.R., Goswami, J., Jun, S.J., Thorne, M., Howell, M., Pusack, T., Kawahara, G., Steffen, L.S., Galdzicki, M., and Kunkel, L.M. (2009). Genetic isolation and characterization of a splicing mutant of zebrafish dystrophin. Human Molecular Genetics 18, 202–211.

Hecht, J.H., Weiner, J.A., Post, S.R., and Chun, J. (1996). Ventricular zone gene-1 (vzg-1) encodes a lysophosphatidic acid receptor expressed in neurogenic regions of the developing cerebral cortex. The Journal of cell biology 135, 1071–1083.

Ho, A.T.V., Palla, A.R., Blake, M.R., Yucel, N.D., Wang, Y.X., Magnusson, K.E.G., Holbrook, C.A., Kraft, P.E., Delp, S.L., and Blau, H.M. (2017). Prostaglandin E2 is essential for efficacious skeletal muscle stem-cell function, augmenting regeneration and strength. Proceedings of the National Academy of Sciences 114, 6675–6684.

Hu, Y., and Smyth, G.K. (2009). ELDA: Extreme limiting dilution analysis for comparing depleted and enriched populations in stem cell and other assays. Journal of Immunological Methods 347, 70–78.

Huxley, H.E. (1963). Electron microscope studies on the structure of natural and synthetic protein filaments from striated muscle. Journal of Molecular Biology 7, 281–IN230.

Ishii, I., Fukushima, N., Ye, X., and Chun, J. (2004). Lysophospholipid receptors: signaling and biology. Annual review of biochemistry 73, 321–354.

Ju, B., Chong, S.W., He, J., Wang, X., Xu, Y., Wan, H., Tong, Y., Yan, T., Korzh, V., and Gong, Z. (2003). Recapitulation of fast skeletal muscle development in zebrafish by transgenic expression of GFP under the mylz2 promoter. Dev Dyn 227, 14–26.

Kim, J.H., and Adelstein, R.S. (2011). LPA(1) -induced migration requires nonmuscle myosin II light chain phosphorylation in breast cancer cells. Journal of cellular physiology 226, 2881–2893.

Konigsberg, U.R., Lipton, B.H., and Konigsberg, I.R. (1975). The regenerative response of single mature muscle fibers isolated in vitro. Developmental biology 45, 260–275.

Kuang, S., Kuroda, K., Le Grand, F., and Rudnicki, M.A. (2007). Asymmetric Self-Renewal and Commitment of Satellite Stem Cells in Muscle. Cell 129, 999–1010.

Lahvic, J.L., Ammerman, M., Li, P., Blair, M.C., Stillman, E.R., Fast, E.M., Robertson, A.L., Christodoulou, C., Perlin, J.R., Yang, S., et al. (2018). Specific oxylipins enhance vertebrate hematopoiesis via the receptor GPR132. Proceedings of the National Academy of Sciences 115, 9252–9257.

Le Grand, F., Jones, A.E., Seale, V., Scime, A., and Rudnicki, M.A. (2009). Wnt7a activates the planar cell polarity pathway to drive the symmetric expansion of satellite stem cells. Cell stem cell 4, 535–547.

Lean, G., Halloran, M.W., Marescal, O., Jamet, S., Lumb, J.-P., and Crist, C. (2019). Ex vivo Expansion of Skeletal Muscle Stem Cells with a Novel Small Compound Inhibitor of eIF2α Dephosphorylation. Regenerative Medicine Frontiers 1, e190003.

Li, P., Lahvic, J.L., Binder, V., Pugach, E.K., Riley, E.B., Tamplin, O.J., Panigrahy, D., Bowman, T.V., Barrett, F.G., Heffner, G.C., et al. (2015). Epoxyeicosatrienoic acids enhance embryonic haematopoiesis and adult marrow engraftment. Nature 523, 468–471.

Liantonio, A., Giannuzzi, V., Picollo, A., Babini, E., Pusch, M., and Conte Camerino, D. (2007). Niflumic acid inhibits chloride conductance of rat skeletal muscle by directly inhibiting the CLC-1 channel and by increasing intracellular calcium. British journal of pharmacology 150, 235–247.

Maesner, C.C., Almada, A.E., and Wagers, A.J. (2016). Established cell surface markers efficiently isolate highly overlapping populations of skeletal muscle satellite cells by fluorescence-activated cell sorting. Skeletal Muscle 6.

Mauro, A. (1961). Satellite cell of skeletal muscle fibers. The Journal of biophysical and biochemical cytology 9, 493–495.

Michalczyk, A., Budkowska, M., Dołęgowska, B., Chlubek, D., and Safranow, K. (2017). Lysophosphatidic acid plasma concentrations in healthy subjects: circadian rhythm and associations with demographic, anthropometric and biochemical parameters. Lipids in health and disease 16, 140–140.

Millay, D.P., O’Rourke, J.R., Sutherland, L.B., Bezprozvannaya, S., Shelton, J.M., Bassel-Duby, R., and Olson, E.N. (2013). Myomaker is a membrane activator of myoblast fusion and muscle formation. Nature 499, 301–305.

Montarras, D., Morgan, J., Collins, C., Relaix, F., Zaffran, S., Cumano, A., Partridge, T., and Buckingham, M. (2005). Direct isolation of satellite cells for skeletal muscle regeneration. Science (New York, NY) 309, 2064–2067.

Moore, J.C., Tang, Q., Yordan, N.T., Moore, F.E., Garcia, E.G., Lobbardi, R., Ramakrishnan, A., Marvin, D.L., Anselmo, A., Sadreyev, R.I., et al. (2016). Single-cell imaging of normal and malignant cell engraftment into optically clear prkdc-null SCID zebrafish. J Exp Med 213, 2575–2589.

Nierobisz, L.S., Cheatham, B., Buehrer, B.M., and Sexton, J.Z. (2013). High-content screening of human primary muscle satellite cells for new therapies for muscular atrophy/dystrophy. Current chemical genomics and translational medicine 7, 21–29.

Oh, J., Sinha, I., Tan, K.Y., Rosner, B., Dreyfuss, J.M., Gjata, O., Tran, P., Shoelson, S.E., and Wagers, A.J. (2016). Age-associated NF-kappaB signaling in myofibers alters the satellite cell niche and re-strains muscle stem cell function. Aging 8, 2871–2896.

Okazaki, K., and Holtzer, H. (1966). Myogenesis: fusion, myosin synthesis, and the mitotic cycle. Proceedings of the National Academy of Sciences of the United States of America 56, 1484–1490.

Partridge, T.A., Grounds, M., and Sloper, J.C. (1978). Evidence of fusion between host and donor myoblasts in skeletal muscle grafts. Nature 273, 306–308.

Poronnik, P., Ward, M.C., and Cook, D.I. (1992). Intracellular Ca2+ release by flufenamic acid and other blockers of the non-selective cation channel. FEBS letters 296, 245–248.

Price, F.D., von Maltzahn, J., Bentzinger, C.F., Dumont, N.A., Yin, H., Chang, N.C., Wilson, D.H., Frenette, J., and Rudnicki, M.A. (2014). Inhibition of JAK-STAT signaling stimulates adult satellite cell function. Nature medicine 20, 1174–1181.

Quinn, M.E., Goh, Q., Kurosaka, M., Gamage, D.G., Petrany, M.J., Prasad, V., and Millay, D.P. (2017). Myomerger induces fusion of non-fusogenic cells and is required for skeletal muscle development. Nature communications 8, 15665.

Rao, T.S., Lariosa-Willingham, K.D., Lin, F.F., Palfreyman, E.L., Yu, N., Chun, J., and Webb, M. (2003). Pharmacological characterization of lysophospholipid receptor signal transduction pathways in rat cerebrocortical astrocytes. Brain Res 990, 182–194.

Richardson, R.M., Pridgen, B.C., Haribabu, B., and Snyderman, R. (2000). Regulation of the human chemokine receptor CCR1. Cross-regulation by CXCR1 and CXCR2. The Journal of biological chemistry 275, 9201–9208.

Sacco, A., Doyonnas, R., Kraft, P., Vitorovic, S., and Blau, H.M. (2008). Self-renewal and expansion of single transplanted muscle stem cells. Nature 456, 502–506.

Sheng, X., Yung, Y.C., Chen, A., and Chun, J. (2015). Lysophosphatidic acid signalling in development. Development (Cambridge, England) 142, 1390–1395.

Sherwood, R.I., Christensen, J.L., Conboy, I.M., Conboy, M.J., Rando, T.A., Weissman, I.L., and Wagers, A.J. (2004). Isolation of Adult Mouse Myogenic Progenitors: Functional Heterogeneity of Cells within and Engrafting Skeletal Muscle. Cell 119, 543–554.

Sinha, M., Jang, Y.C., Oh, J., Khong, D., Wu, E.Y., Manohar, R., Miller, C., Regalado, S.G., Loffredo, F.S., Pancoast, J.R., et al. (2014). Restoring systemic GDF11 levels reverses age-related dysfunction in mouse skeletal muscle. Science (New York, NY) 344, 649–652.

Stapleton, C.M., Mashek, D.G., Wang, S., Nagle, C.A., Cline, G.W., Thuillier, P., Leesnitzer, L.M., Li, L.O., Stimmel, J.B., Shulman, G.I., et al. (2011). Lysophosphatidic Acid Activates Peroxisome Proliferator Activated Receptor-γ in CHO Cells That Over-Express Glycerol 3-Phosphate Acyltransferase-1. PLOS ONE 6, e18932.

Tabebordbar, M., Wang, E.T., and Wagers, A.J. (2013). Skeletal Muscle Degenerative Diseases and Strategies for Therapeutic Muscle Repair. In Annu Rev Pathol Mech Dis, pp. 441–475.

Tanaka, K.K., Hall, J.K., Troy, A.A., Cornelison, D.D., Majka, S.M., and Olwin, B.B. (2009). Syndecan-4-expressing muscle progenitor cells in the SP engraft as satellite cells during muscle regeneration. Cell stem cell 4, 217–225.

Tierney, M.T., Aydogdu, T., Sala, D., Malecova, B., Gatto, S., Puri, P.L., Latella, L., and Sacco, A. (2014). STAT3 signaling controls satellite cell expansion and skeletal muscle repair. Nature medicine 20, 1182–1186.

Westerfield, M. (2007). The Zebrafish Book. A Guide for the Laboratory Use of Zebrafish (Danio rerio) (University of Oregon Press: Eugene).

Xu, C., Tabebordbar, M., Iovino, S., Ciarlo, C., Liu, J., Castiglioni, A., Price, E., Liu, M., Barton, E.R., Kahn, C.R., et al. (2013). A zebrafish embryo culture system defines factors that promote vertebrate myogenesis across species. Cell 155, 909–921.

Xu, Y.-J., Tappia, P.S., Goyal, R.K., and Dhalla, N.S. (2008). Mechanisms of the lysophosphatidic acid-induced increase in [Ca(2+)](i) in skeletal muscle cells. Journal of cellular and molecular medicine 12, 942–954.

Ye, X., and Chun, J. (2010). Lysophosphatidic acid (LPA) signaling in vertebrate reproduction. Trends in endocrinology and metabolism: TEM 21, 17–24.

Yin, H., Price, F., and Rudnicki, M.A. (2013). Satellite cells and the muscle stem cell niche. Physiological Reviews 93, 23–67.

